# A conserved lncRNA regulates trehalose-glucose homeostasis through direct RNA-RNA interactions

**DOI:** 10.64898/2026.07.14.738560

**Authors:** Vikram Nichit, Deepti Wagh, Anand Kumar Shukla, Narendra Kadoo, Rakesh Joshi

## Abstract

Trehalose is a primary circulating sugar in insects and essential for energy homeostasis, yet its non-coding RNA-based regulatory circuitry remains enigmatic. Here, we characterize a conserved long non-coding RNA, lncRNA1, as a post-transcriptional regulator of trehalose-glucose homeostasis in Lepidoptera. lncRNA1 encodes a structurally stable, pseudoknot-containing transcript that is strongly induced upon trehalose pathway perturbation and exhibits a reciprocal developmental expression pattern relative to the trehalose metabolism enzymes. RNAi-mediated silencing of lncRNA1 in *Helicoverpa armigera* elevates TPS/TPP and Treh transcript abundance, increases enzyme activities, reduces haemolymph trehalose, raises glucose. This drives broad transcriptomic and metabolomic reprogramming of carbohydrate, lipid, and growth-signalling pathways, resulting in accelerated larval growth. Overexpression of lncRNA1 reverses these phenotypes. Mechanistically, lncRNA1 physically associates with TPS/TPP and Treh mRNAs through evolutionarily conserved sequence motifs, modulating their post-transcriptional dynamics. Targeted deletion of these motifs abolishes regulatory activity and disrupts metabolic homeostasis. This regulatory axis is functionally conserved in *Spodoptera frugiperda*, validated across loss-of-function, gain-of-function, and cell-based systems. Our findings reveal a conserved lncRNA-based layer of post-transcriptional control over insect energy metabolism.

**Graphical Abstract:** 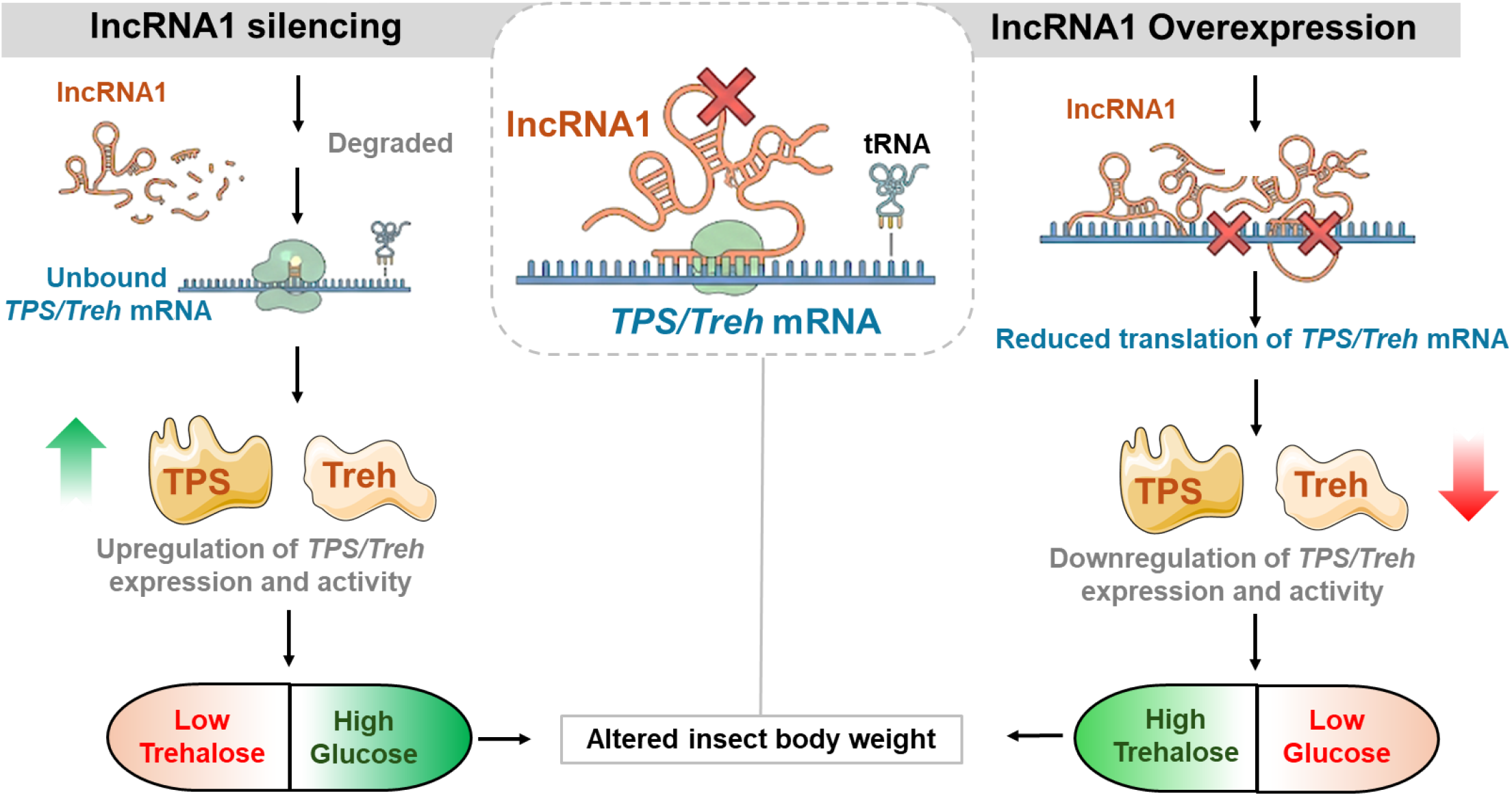

## Introduction

Trehalose metabolism is a crucial biochemical pathway in insects, involving both anabolic and catabolic processes essential for maintaining energy homeostasis and enabling adaptation to environmental stresses such as desiccation, starvation, and cold stress ^1–4^. Trehalose serves as the primary circulating sugar in insect hemolymph and functions as a vital energy source ^5,6^. Its synthesized in insect fat body by trehalose 6-phosphate synthase/phosphatase (*TPS/TPP*), while its hydrolysed into glucose by trehalase (*Treh*) for energy production ^7^. Trehalose regulates energy homeostasis during development, flight and stress recovery ^6,8,9^.

Trehalose metabolism is regulated spatiotemporally at multiple levels. Several transcription factors (TFs) involved in insulin signaling and metamorphosis directly regulate trehalose metabolic genes ^9^. Forkhead box O (FoxO) has been shown to bind to the *TPS* promoter in *Aedes aegypti* and *Tribolium castaneum* to promotes trehalose synthesis through activation of gluconeogenesis and glycogenolysis ^10^. In *Drosophila*, the E2F1–Dp complex regulates *TPS* expression in the fat body, while Kruppel (Kr) acts as a positive regulator of *TPS* in *A. aegypti* ^11,12^. In contrast, the ecdysone receptor (EcR) and its downstream effector *E74A* repress *TPS*, linking hormonal cues to trehalose regulation. Trehalose transport and catabolism are also transcriptionally controlled; TGF-β signaling induces trehalose transporter (*TRET-1*) expression in the *D. melanogaster* blood–brain barrier ^13^. Despite a considerable understanding of these regulatory mechanisms, the fine-scale regulation of trehalose metabolism, particularly the role of non-coding RNAs in this process, remains largely unexplored.

Long non-coding RNAs (lncRNAs) are transcripts longer than 200 nucleotides that lack protein-coding potential ^14,15^ and have emerged as important regulators of gene expression across diverse organisms, including insects ^16^. In insect systems, lncRNAs have been implicated in a wide range of biological processes, including development, metabolism and reproduction ^17,18^. Several lncRNAs have been functionally characterised in ecdysteroid signalling-associated developmental processes, such as *iab*, which regulates Hox gene expression during segmentation, and *acal*, which modulates the JNK signalling pathway ^19,20^. At the same time, CR33938 contributes to appendage development ^21^. In metabolic contexts, IBIN and IRAR have been linked to carbohydrate metabolism and nutrient-responsive insulin signalling, respectively ^22,23^. lncRNAs also participate in reproductive regulation, with *msa*, *iab-8* and roX RNAs (*roX1* and *roX2*) implicated in fertility, dosage compensation and gene regulation, and others contributing to spermatogenesis and oogenesis ^24–27^. Furthermore, lncRNAs such as *yar* have been associated with circadian regulation ^28^. Despite these advances, mechanistic insights into lncRNA-mediated regulation in insects remain limited, particularly with respect to core metabolic pathways such as trehalose metabolism. Understanding lncRNA-mediated control of trehalose metabolism is especially relevant in agricultural pests such as *Helicoverpa armigera* and *Spodoptera frugiperda*, whose development and survival are tightly linked to carbohydrate homeostasis.

Here, we integrated computational prediction, transcriptomic and metabolomic profiling, transcriptional stability analysis, and functional assays to investigate the role of long non-coding RNAs (lncRNAs) in regulating trehalose metabolism in lepidopteran insects. We identified a structurally stable and evolutionarily conserved lncRNA, designated *lncRNA1*, that responded to perturbations of the trehalose metabolic pathway. The functional significance of *lncRNA1* was examined through RNA interference-mediated silencing and overexpression approaches in *Helicoverpa armigera* and *Spodoptera frugiperda*. To further elucidate its mode of action, we assessed the transcriptional stability of *lncRNA1* and its associated target genes using Actinomycin D treatment, revealing coordinated expression dynamics consistent with a regulatory role at the post-transcriptional level. We also evaluated potential *lncRNA1*–mRNA interactions with key trehalose metabolic transcripts and investigated the contribution of conserved sequence regions to its regulatory activity. Collectively, these analyses provide comprehensive evidence that a conserved lncRNA participates in the post-transcriptional modulation of trehalose metabolism and metabolic homeostasis in lepidopteran insects.

## Results

### *HalncRNA1* with stable loop structure formation showed elevated expression on trehalose metabolism inhibition

As trehalose metabolism is tightly regulated in spatio-temporal manner, hence, to understand potential lncRNA diversity and their expression dynamics upon trehalose metabolism modulation global lncRNA prediction and expression analysis was performed. A stepwise computational pipeline integrating annotation-based filtering, RNAfold stability assessment, coding potential evaluation (lncRNA_Mdeep), and pseudoknot prediction (IPknot++) refined 1733 annotated ncRNAs to 185 high-confidence lncRNAs exhibiting stable secondary structures with predicted pseudoknots (**Fig. 1A and Supplementary Data 2)**. Out of 1,663 annotated lncRNAs analyzed for thermodynamic stability, 283 transcripts clustered within the fourth quadrant of the two-dimensional MFE_norm versus Diversity_norm stability framework, indicating strong structural stability characterized by highly negative normalized free energy and low conformational variability (**Fig. 1B and Supplementary Data 2)**.

**Fig 1.**
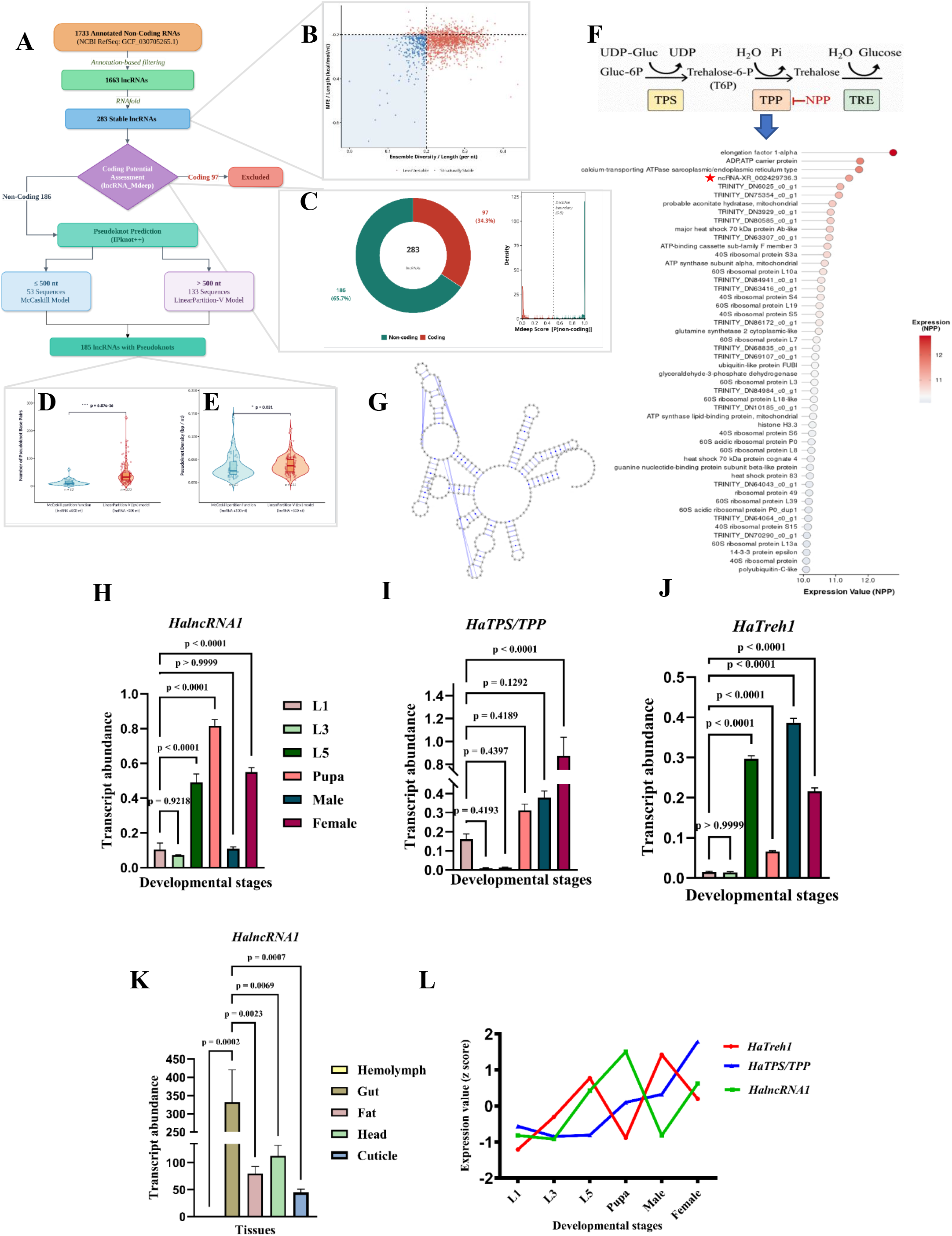
Integrated expression profiling of *HalncRNA1* and trehalose metabolism genes across developmental stages, tissues, and trehalose metabolism inhibition. (A) Integrated computational workflow for stability-guided identification and pseudoknot characterization of lncRNAs in *H armigera.* (B) Scatter plot of normalized ensemble diversity (per nt) versus normalized MFE (kcal/mol/nt) for 1,663 lncRNAs, highlighting 283 transcripts clustered in the fourth quadrant representing structurally stable candidates with low diversity and highly negative free energy. **(C)** Coding potential classification of 283 structurally stable lncRNAs using lncRNA Mdeep, identifying 186 transcripts as non-coding (probability ≥ 0.5) while excluding 97 predicted coding transcripts. **(D)** Comparison of the number of pseudoknot base pairs between short (≤500 nt; McCaskill partition function) and long (>500 nt; LinearPartition-V model) lncRNAs among 186 validated non-coding candidates, showing a highly significant difference (Wilcoxon rank–sum test, p = 6.87 × 10⁻¹⁶). **(E)** Comparison of pseudoknot density (bp/nt) between short (≤500 nt) and long (>500 nt) lncRNAs, indicating a significant difference (Wilcoxon rank–sum test, p = 0.031) and suggesting length-dependent variation in structural complexity. **(F)** Schematic representation of trehalose metabolism showing inhibition of trehalose-6-phosphate phosphatase (TPP) by N-(phenylthio) phthalimide (NPP). Downstream transcriptomic analysis highlights differentially expressed genes (DEGs) in response to pathway disruption. The dot plot represents normalized expression values of DEGs under NPP treatment, with color intensity indicating expression levels. The lncRNA *XR_002429736.3* (*HalncRNA1*) is marked (red star) among the top differentially expressed transcripts. **(G)** Predicted secondary structure of *HalncRNA1*, highlighting pseudoknot interactions. Quantitative reverse-transcriptase PCR (qRT-PCR) analysis showing transcript abundance of (H) *HalncRNA1*, (I) *HaTPS/TPP*, and (J) *HaTreh1* across developmental stages, including first instar larvae (L1), third instar larvae (L3), fifth instar larvae (L5), pupa (P), male (M), and female (F). **(K)** Tissue-specific expression patterns of *HalncRNA1*, in fifth instar larvae, including hemolymph, fat body, gut, head, and cuticle. **(L)** Correlation analysis between *HalncRNA1*, *HaTPS/TPP and HaTreh1*, expression levels. Differences among developmental stages were assessed using one-way analysis of variance (ANOVA), as appropriate. Statistical significance was determined at *P* < 0.05, with exact *P*-values indicated in the figure.

Coding potential assessment using lncRNA_Mdeep further refined this set, classifying 186 transcripts as non-coding (probability ≥ 0.5), while 97 transcripts predicted as coding were excluded from downstream analysis (**Fig. 1C and Supplementary Data 2)**. Among the 186 validated non-coding lncRNAs, 185 were predicted to contain pseudoknot structures. Comparative analysis of pseudoknot features between transcripts analyzed using the McCaskill partition function (≤500 nt) and the LinearPartition-V (lpv) model (>500 nt) revealed a highly significant difference in the number of pseudoknot base pairs (p = 6.87 × 10⁻¹⁶) and a significant difference in pseudoknot density (bp/nt) (p = 0.031), indicating length-dependent structural complexity among stable lncRNA candidates (**Fig. 1D, E and Supplementary Data 2)**.

*De novo* transcriptomic analysis of *H. armigera* following treatment with N-(phenylthio) phthalimide (NPP) inhibitor of *Ha*TPP, revealed significant transcriptional alterations ^29^. Interestingly, among the identified transcripts, ncRNA-XR002429736.3, designated as *HalncRNA1*, ranked among the top ten differentially expressed genes in response to NPP treatment (**Fig. 1F**). Subsequent analysis using the NCBI database and our transcriptomic datasets confirmed that this transcript exhibits high structural stability, a hallmark of lncRNAs. The *HalncRNA1* (298 nt) exhibited strong thermodynamic stability with a minimum free energy (MFE) of −84.70 kcal/mol, ensemble free energy of −88.20 kcal/mol, and centroid free energy of −82.20 kcal/mol **(Supplementary Data 2)**. The MFE frequency was 0.34%, indicating that although the minimum energy structure is thermodynamically favorable, alternative conformations contribute to the structural ensemble. The ensemble diversity value of 44.27 suggests moderate conformational variability. The predicted secondary structure of *HalncRNA1*, highlighting pseudoknot interactions. The transcript (298 nt) contains eight pseudoknot base pairs, corresponding to a pseudoknot density of 0.0268 bp/nt, indicating the presence of complex higher-order structural features (**Fig. 1G and Supplementary Data 2)**. Coding potential analysis using lncRNA_Mdeep classified *lncRNA1* as non-coding with extremely high confidence (P = 0.9999996) **(Supplementary Data 2)**. Transcript classification using gffcompare ^30^ assigned class code “=” indicating an exact match to the reference annotation, corresponding to locus LOC110379414 on the positive strand. Genomic context analysis confirmed that the transcript is located in an intergenic region between LOC135116975 and LOC110379411, without overlapping any protein-coding genes, and thus classifies it as a long intergenic non-coding RNA (lincRNA).

### *HalncRNA1* showed a reciprocal expression pattern compared to trehalose metabolism genes

To unravel the correlation in expression patterns between *HalncRNA1* and genes involved in trehalose metabolism, we performed spatiotemporal expression analysis. Developmental-stages-wide expression dynamics of *HalncRNA1* revealed high expression in the fifth instar larva (L5) and the pupal stage, a critical developmental phase associated with extensive metabolic remodeling. Notably, *HalncRNA1* expression was higher in female adults compared to male adults (**Fig. 1H**). In contrast, *HaTPS/TPP* expression in L5 was found to be relatively lower compared to early instars, while pupal expression was higher (**Fig. 1I**). Furthermore, *HaTreh1* exhibited elevated expression during the L5 stage but was markedly reduced in the pupal stage. Consistent with this opposing trend, *HaTreh1* showed higher transcript abundance in male adults relative to females (**Fig. 1J**). Tissue-specific expression analysis further revealed that *HalncRNA1* was predominantly expressed in the gut, head and fat body, the major sites of energy reserve mobilization, utilization, and carbohydrate metabolism in insects (**Fig. 1K**). *HaTPS/TPP*, *HaTreh1*, and *HalncRNA1* exhibited largely reciprocal expression patterns based on z-score normalization (**Fig. 1L**). Notably, *HalncRNA1* expression peaked at the pupal stage and exhibited an inverse relationship with *HaTreh1*, suggesting a potential regulatory role in trehalose metabolism.

### *lncRNA1* silencing alters trehalose accumulation in lepidopteran insects

To investigate the biological function of *HalncRNA1*, RNAi-mediated silencing was performed in *H. armigera* larvae using bacterially expressed dsRNA (**Fig. 2A**). Agarose gel electrophoresis confirmed successful production of *HalncRNA1*-specific dsRNA in HT115 cells carrying the recombinant L4440 vector **(Fig. S1A)**. Phenotypic assessment revealed that *HalncRNA1*-silenced larvae were visibly larger than control larvae and exhibited a progressive increase in average body weight from 3 to 6 days post-treatment (**Fig. 2B, C**). qRT-PCR analysis demonstrated a significant reduction in *HalncRNA1* transcript abundance in ds*HalncRNA1*-treated larvae compared with the control group, confirming effective gene silencing (**Fig. 2D**). Further analysis revealed an evident upregulation of *HaTPS/TPP* (**Fig. 2E**) and *HaTreh1* expressions (**Fig. 2F**), whereas *HaTreh2* expression remained unchanged in *HalncRNA1*-silenced tissues **(Fig. S1B)**. These transcriptional changes were consistent with enzyme activity assays, which showed significantly elevated *Ha*TPP (**Fig. 2G**) and *Ha*Treh1 (**Fig. 2H**) activities.

**Fig 2.**
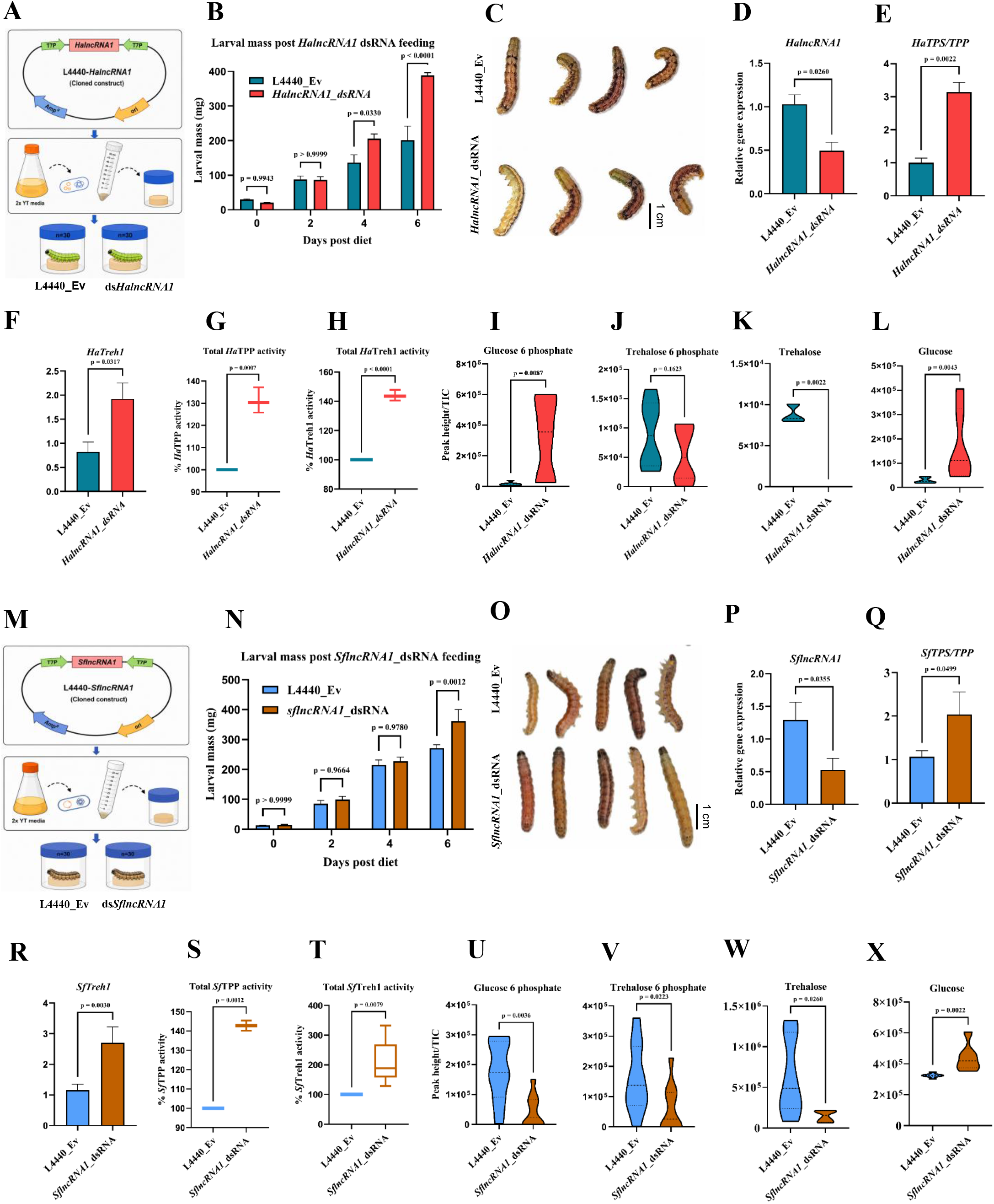
*lncRNA1* silencing alters trehalose accumulation in *H. armigera* and *S. frugiperda* insect. **(A)** Schematic illustration of the RNAi workflow for *HalncRNA1* silencing in H. armigera. The *HalncRNA1* fragment was cloned into the L4440 vector, expressed in HT115 cells, and delivered orally to larvae for functional analysis. **(B)** Larval mass measured over time following dsRNA feeding, indicating increased growth upon *HalncRNA1* silencing. **(C)** Representative images of control (upper panel) and dsHalncRNA1-treated (lower panel) *H. armigera* larvae at 6 days post-treatment. *HalncRNA1*-silenced larvae exhibited increased body size compared with control larvae. Scale bar = 1 cm. **(D)** qRT-PCR analysis showing efficient knockdown of *HalncRNA1* in dsRNA-treated larvae compared to L4440 empty vector (Ev) control. **(E, F)** Relative expression levels of *HaTPS/TPP and HaTreh1* following *HalncRNA1* silencing. **(G, H)** Corresponding enzyme activity assays showing changes in *Ha*TPP and *Ha*Treh1 activity upon dsRNA treatment. **(I–L)** Metabolite profiling showing levels of glucose-6-phosphate, trehalose-6-phosphate, trehalose, and glucose in control and dsRNA-treated larvae. Differences between L4440 Ev and *HalncRNA1* dsRNA treatments were assessed using an unpaired two-tailed Student’s *t*-test. Data are presented as mean ± SEM from three biological replicates. The exact *P*-value is indicated in the figure, with *P* < 0.05 considered statistically significant. **(M)** Schematic illustration of the RNAi workflow for *SflncRNA1* silencing in *S. frugiperda*. The *SflncRNA1* fragment was cloned into the L4440 vector, expressed in HT115 cells, and delivered orally to larvae for functional analysis **(N)** Larval mass measured over time following dsRNA feeding, showing increased growth in *SflncRNA1*-silenced larvae compared to L4440 empty vector (Ev) control. **(O)** Representative images of ds*SflncRNA1*-treated (upper panel) and control (lower panel) *S.frugiperda* larvae at 6 days post-treatment. *SflncRNA1*-silenced larvae exhibited increased body size compared with control larvae. Scale bar = 1 cm. **(P)** qRT-PCR validation of *SflncRNA1* knockdown in dsRNA-treated samples. **(Q, R)** Relative expression levels of *SfTPS/TPP* and *SfTreh1* following *SflncRNA1* silencing. **(S, T)** Corresponding enzyme activity assays showing increased *Sf*TPP and *Sf*Treh1 activity upon dsRNA treatment. **(U–X)** Metabolite profiling showing levels of glucose-6-phosphate, trehalose-6-phosphate, trehalose, and glucose in control and dsRNA-treated larvae. Differences between L4440 Ev and *SflncRNA1* dsRNA treatments were assessed using an unpaired two-tailed Student’s *t*-test. Data are presented as mean ± SEM from three biological replicates. The exact *P*-value is indicated in the figure, with *P* < 0.05 considered statistically significant.

Moreover, metabolomic profiling of the silenced larvae revealed a substantial decrease in trehalose content, accompanied by a corresponding increase in glucose levels, indicative of a metabolic shift toward enhanced trehalose catabolism (**Fig. 2I–L**). Additionally, the *HaST9, HaST29, HaST46*, *and HaST64* sugar transporters exhibited dysregulated expression in *HalncRNA1*-silenced tissues **(Fig. S1C-F)**. Key TFs associated with glucose metabolism, namely *FOXO* and *IGF2*, were downregulated and upregulated, respectively **(Fig. S1G, H).** Notably, the developmental regulator *HaAL* and the cell cycle–associated transcription factor *HaE2F* were significantly upregulated, while the expression levels of the remaining transcription factors showed no significant changes. **(Fig. S1I-O)**.

To validate the conservation of *lncRNA1’s* role to regulate trehalose metabolism in *S. frugiperda*, we performed *lncRNA1* silencing (**Fig. 2M**). *SflncRNA1* was identified based on sequence similarity search using *HalncRNA1* as a query against the *S. frugiperda* Refseq database. The top hit identified from this analysis was (XR_004783399.2), having 90.43 % sequence identity (Query cover: 58%) with *HalncRNA1*.

Likewise, *HalncRNA1*, phenotypic assessment revealed that *SflncRNA1*-silenced larvae were visibly larger than control larvae and exhibited a progressive increase in average body weight on day 6 post-treatment (**Fig. 2N, O**). *SflncRNA1* silencing resulted in a significant reduction in its transcript levels (**Fig. 2P**). Expression analysis revealed an evident upregulation of *SfTPS/TPP* (**Fig. 2Q**) and *SfTreh1* (**Fig. 2R**). These transcriptional changes were consistent with enzyme activity assays, which showed significantly elevated *Sf*TPP (**Fig. 2S**) and *Sf*Treh1 (**Fig. 2T**) activities. Moreover, metabolomic profiling of *SflncRNA1*-silenced larvae revealed changes consistent with those observed upon *HalncRNA1* silencing, characterized by decreased trehalose levels and a concomitant increase in glucose, indicating enhanced trehalose catabolism. (**Fig. 2U–X**). Collectively, these findings suggest that silencing of *lncRNA1* triggers the upregulation of *TPS/TPP* and *Treh* transcripts, thereby perturbing trehalose and glucose homeostasis and leading to altering carbohydrate metabolism.

### *HalncRNA1* silencing reprograms trehalose and glucose metabolism, driving a shift in carbohydrate utilization pathways

To elucidate the molecular pathways regulated by *HalncRNA1*, transcriptome profiling was conducted in whole insects following RNAi-mediated silencing of *HalncRNA1*. Differential expression analysis identified widespread changes in gene expression, revealing extensive transcriptional reprogramming between ds*HalncRNA1*-treated and control insects (**Fig. 3A and Fig. S2A)**. Functional annotation of the differentially expressed genes indicated significant modulation of pathways related to energy metabolism, nutrient utilization, hormonal regulation, and developmental processes, highlighting the central role of *HalncRNA1* in maintaining metabolic and physiological homeostasis. Notably, several genes involved in energy metabolism and nutrient utilization, including putative ATP synthase subunit f, phosphoglyceromutase, β-hydroxyacid dehydrogenase 1, D-erythronate dehydrogenase, and adenylate kinase isoenzyme 5, were markedly induced, suggesting enhanced metabolic activity and energy turnover. Genes associated with hormonal regulation and developmental processes, such as juvenile hormone epoxide hydrolase-like, β-1,4-N-acetylgalactosaminyltransferase bre-4, and argininosuccinate synthase, were also significantly upregulated. Furthermore, increased expression of mitochondrial enolase and aldo-keto reductase family 1 member B1 indicates activation of mitochondrial metabolism and redox-balancing mechanisms.

**Fig 3.**
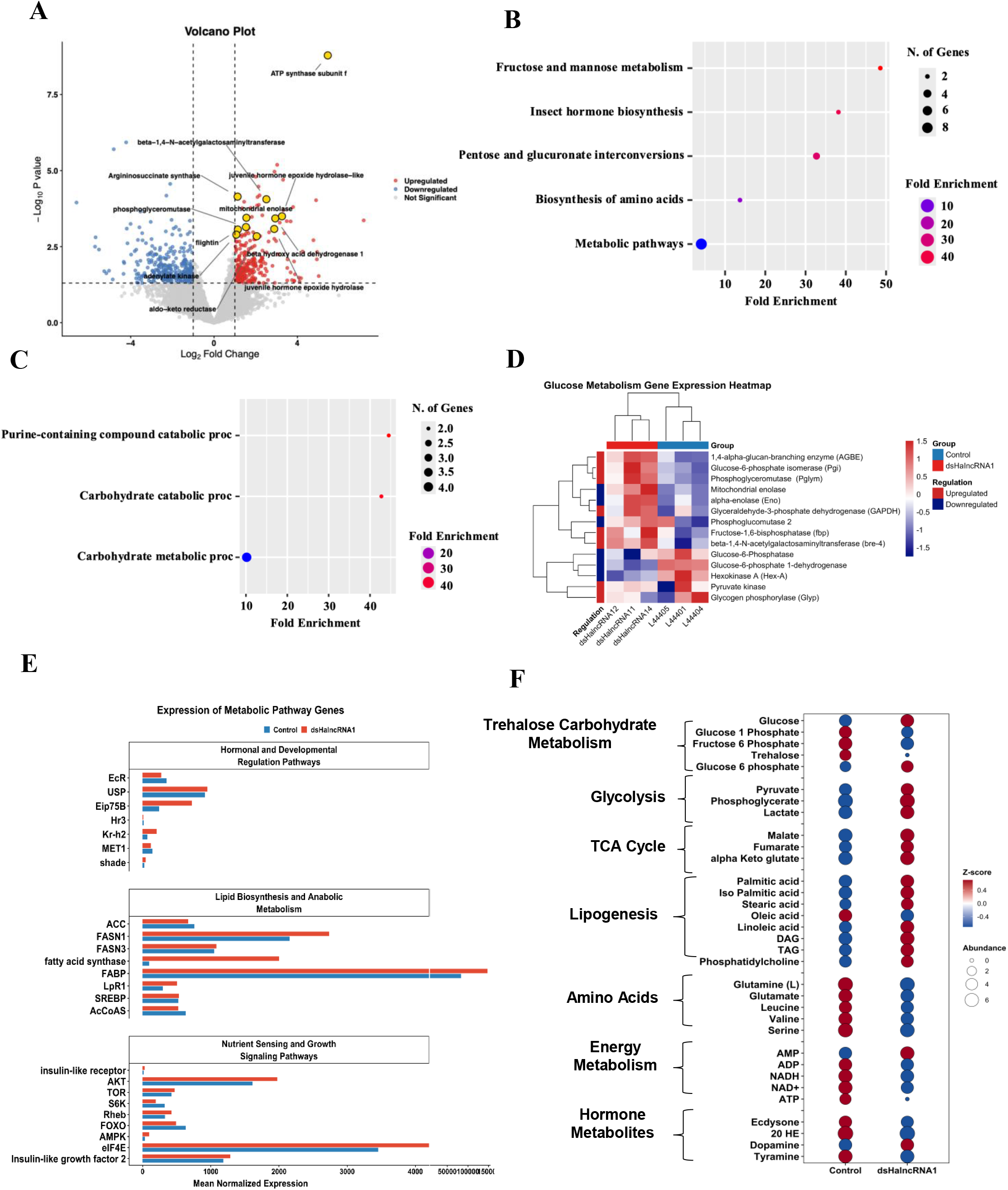
*HalncRNA1* silencing reprograms trehalose and glucose metabolism, driving a shift in carbohydrate utilization pathways. **(A)** Differential gene expression analysis of *HalncRNA1*-silenced insects. Volcano plot showing differentially expressed genes (DEGs) between ds*HalncRNA1*-treated and control (L4440) insects. The x-axis represents the log₂ fold change (Log_2F_C), while the y-axis represents the −log_10(_P-value). Red dots indicate significantly upregulated genes, blue dots indicate significantly downregulated genes, and gray dots represent non-significant transcripts. Selected significantly regulated genes involved in energy metabolism, carbohydrate metabolism, hormonal regulation, and growth-associated genes are highlighted. **(B)** KEGG pathway enrichment analysis highlighting significantly enriched pathways, including fructose and mannose metabolism, insect hormone biosynthesis, pentose and glucuronate interconversions, and amino acid biosynthesis. **(C)** Gene Ontology (GO) biological process enrichment analysis showing enrichment of carbohydrate metabolic and catabolic processes, along with purine-containing compound catabolic processes. Dot size represents the number of genes, and color indicates fold enrichment. **(D)** Expression profiling of glucose and glycogen metabolism-associated genes following *HalncRNA1* silencing in *Helicoverpa armigera*. Heatmap showing the relative expression patterns of genes involved in carbohydrate metabolism in control (L4440) and ds*HalncRNA1*-treated insects. Expression values were normalized and Z-score transformed prior to hierarchical clustering. Red and blue colors indicate relatively higher and lower expression levels, respectively. **(E)** Expression profiles of genes associated with nutrient sensing, lipid biosynthesis, and hormonal signaling pathways following *HalncRNA1* silencing. Bar plots represent normalized transcript abundance of selected genes in control (L4440) and ds*HalncRNA1*-treated insects. Genes were grouped into three functional categories: nutrient sensing and growth signaling pathways (insulin-like receptor, PI3K/AKT/TOR signaling components, AMPK, eIF4E, and insulin-like growth factor 2), lipid biosynthesis and anabolic metabolism (ACC, FASN1, FASN3, fatty acid synthase, FABP, LpR1, SREBP, and AcCoAS), and hormonal and developmental regulation pathways (EcR, USP, Eip75B, Hr3, Kr-h2, MET1, and shade). **(F)** Metabolic reprogramming associated with *HalncRNA1* silencing in *Helicoverpa armigera*. Bubble plot showing the relative abundance of key metabolites involved in carbohydrate metabolism, glycolysis, the tricarboxylic acid (TCA) cycle, lipogenesis, amino acid metabolism, energy metabolism, and hormonal regulation in control and ds*HalncRNA1*-treated insects. Bubble color represents the Z-score-normalized abundance of each metabolite, with red and blue indicating relatively higher and lower abundance, respectively, while bubble size corresponds to metabolite abundance.

Pathway enrichment analysis revealed significant overrepresentation of carbohydrate-related metabolic pathways following *HalncRNA1* silencing (**Fig. 3B**). Fructose and mannose metabolism showed the highest fold enrichment, followed by pentose and glucuronate interconversions. Gene ontology (GO) analysis further highlighted enrichment of carbohydrate metabolic and catabolic processes (**Fig. 3C**). Normalized expression analysis of trehalose metabolism and sugar transporter genes from RNASeq data revealed distinct transcriptional changes upon *HalncRNA1* silencing **(Fig. S2B)**. The expression of *HaTPS/TPP*, *HaTreh1*, and the sugar transporter gene *HaST69* was markedly upregulated in ds*HalncRNA1*-treated samples compared to the L4440 control, consistent with qRT-PCR validation. In contrast, *HaTreh2* and the sugar transporter genes *HaST9, HaST29, HaST46,* and *HaST64* were downregulated following *HalncRNA1* silencing. Silencing of *HalncRNA1* resulted in pronounced transcriptional reprogramming of genes associated with glucose and glycogen metabolism (**Fig. 3D**). Several genes involved in glycogen synthesis and carbohydrate turnover, including 1,4-alpha-glucan-branching enzyme (AGBE), glucose-6-phosphate isomerase (Pgi), glycogen synthase (Pglym), glyceraldehyde-3-phosphate dehydrogenase (GAPDH), fructose-1,6-bisphosphatase (fbp), beta-1,4-N-acetylgalactosaminyltransferase (bre-4), pyruvate kinase, and glycogen phosphorylase (Glyp), were upregulated following *HalncRNA1* silencing. In contrast, phosphoglucomutase, glucose-6-phosphatase, glucose-6-phosphate 1-dehydrogenase, and hexokinase A were downregulated relative to the controls. Collectively, these transcriptional changes indicate a major reorganization of carbohydrate metabolism in response to *HalncRNA1* silencing. The simultaneous upregulation of glycogen phosphorylase and pyruvate kinase suggests enhanced glycogen mobilization and carbon flux toward energy-generating pathways, while the altered expression of key glycolytic and glucose-processing enzymes reflects a rewiring of glucose utilization.

Silencing of *HalncRNA1* resulted in extensive transcriptional changes in genes associated with nutrient sensing, lipid metabolism, and hormonal signaling pathways (**Fig. 3E, Supplementary Data 3)**. Notably, genes involved in lipid biosynthesis and anabolic metabolism, including FASN1, FASN3, fatty acid synthase, FABP, LpR1, and eIF4E, were upregulated, indicating enhanced fatty acid synthesis, lipid transport, and protein translation. In addition, the increased expression of AKT, Rheb, AMPK, and insulin-like growth factor 2 (IGF2) suggests activation of growth-promoting pathways, whereas components of hormonal signaling, including EcR, FOXO, MET1, and shade, were downregulated.

Metabolomic profiling revealed extensive metabolic reprogramming following *HalncRNA1* silencing (**Fig. 3F, Supplementary Data 3)**. Compared with the control, *dsHalncRNA1*-treated insects exhibited elevated levels of glucose, glucose-6-phosphate, pyruvate, lactate, malate, fumarate, α-ketoglutarate, and several lipid-associated metabolites including palmitic acid, linoleic acid, DAG, TAG, and phosphatidylcholine, indicating enhanced carbohydrate catabolism, TCA cycle activity, and lipid biosynthesis. In contrast, trehalose, multiple amino acids (glutamine, glutamate, leucine, valine, and serine), and the hormones ecdysone, 20-hydroxyecdysone, and tyramine were reduced, whereas dopamine accumulated in silenced insects.

Collectively, the transcriptomic and metabolomic analyses demonstrate that *HalncRNA1* functions as a key regulator of carbohydrate homeostasis in *H. armigera*. Silencing of *HalncRNA1* promotes trehalose mobilization, glucose accumulation, enhanced glycolytic and TCA cycle activity, and activation of nutrient-sensing and lipogenic pathways, resulting in a metabolic state that favors efficient energy utilization and biomass production. These changes were accompanied by extensive transcriptional reprogramming of genes involved in carbohydrate metabolism, energy turnover, and growth-related processes. Together, these findings provide a mechanistic explanation for the increased body size and body weight observed in *HalncRNA1*-silenced insects.

### *lncRNA1* overexpression suppresses trehalose biosynthesis and degradation pathways

To elucidate the functional role of *lncRNA1* in trehalose metabolism regulation, *HalncRNA1* was transiently overexpressed (OE) in *H. armigera* larvae (**Fig. 4A**). qRT-PCR confirmed significant upregulation of the target *HalncRNA1* (**Fig. 4B**). Expression profiling of trehalose metabolism genes revealed substantial downregulation of *HaTPS/TPP* (**Fig. 4C**), *HaTreh1* (**Fig. 4D),** and *HaTreh2* showed downregulation **(Fig. S3A)** in *HalncRNA1*_OE insects. Consistently, enzymatic assays demonstrated a significant decrease in trehalase activity (**Fig. 4E**). Metabolomic profiling further revealed markedly reduced glucose accumulation compared with control groups (**Fig. 4F-I**).

**Fig 4.**
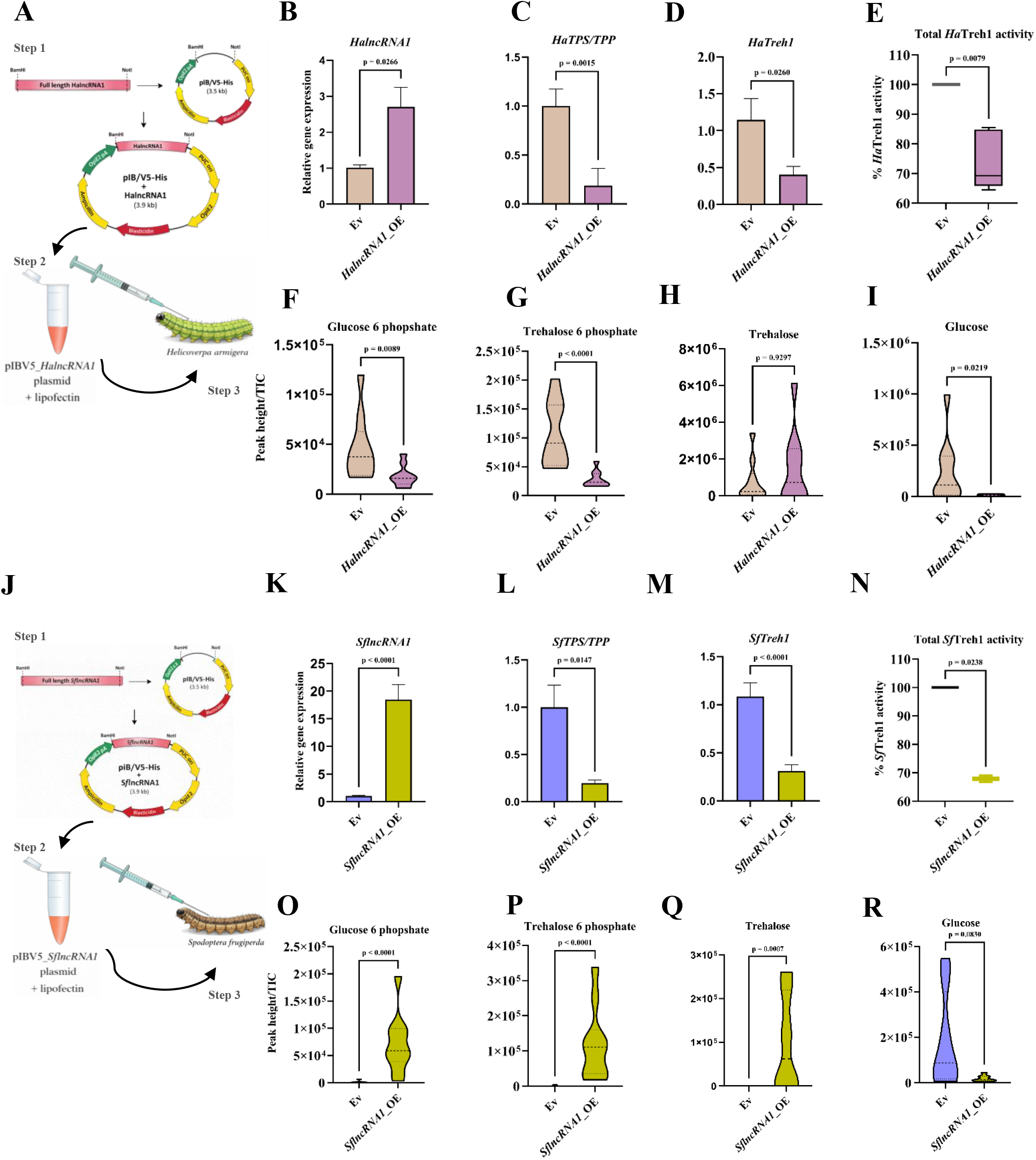
*lncRNA1* overexpression suppresses trehalose biosynthesis and degradation pathways in *H. armigera* and *S. frugiperda.* **(A)** Schematic representation of cloning of full-length *HalncRNA1* into the pIB/V5-His expression vector and subsequent delivery into *H. armigera* larvae using lipofectin-mediated transfection. **(B)** qRT-PCR validation of *HalncRNA1* overexpression (OE) compared to empty vector (Ev) control. **(C, D)** Relative expression levels of *HaTPS/TPP* and *HaTreh1* following *HalncRNA1*_OE. **(E)** Total Treh1 enzyme activity in Ev and OE samples. **(F–I)** Metabolite profiling showing levels of glucose-6-phosphate, trehalose-6-phosphate, trehalose, and glucose in control and *HalncRNA1*-overexpressing insects. Differences between Ev and *HalncRNA1* OE treatments were assessed using an unpaired two-tailed Student’s *t*-test. Data are presented as mean ± SEM from three biological replicates. The exact *P*-value is indicated in the figure, with *P* < 0.05 considered statistically significant. **(J)** Schematic representation of cloning of full-length *SflncRNA1* into the pIB/V5-His expression vector and delivery into *S. frugiperda* larvae using lipofectin-mediated transfection. **(K)** qRT-PCR validation of *SflncRNA1* overexpression (OE) compared to empty vector (Ev) control. **(L, M)** Relative expression levels of *SfTPS/TPP* and *SfTreh1* following *SflncRNA1* overexpression. **(N)** Total Treh1 enzyme activity in Ev and OE samples. **(O–R)** Metabolite profiling showing levels of glucose-6-phosphate, trehalose-6-phosphate, trehalose, and glucose in control and *SflncRNA1*-overexpressing insects. Differences between Ev and *SflncRNA1* OE treatments were assessed using an unpaired two-tailed Student’s *t*-test. Data are presented as mean ± SEM from three biological replicates. The exact *P*-value is indicated in the figure, with *P* < 0.05 considered statistically significant.

Similarly, the functional role of *SflncRNA1* in trehalose metabolism was confirmed by *SflncRNA1* overexpression (**Fig. 4J**). Upon transient overexpression, significant upregulation of *SflncRNA1* was observed in the whole insect body (**Fig. 4K**). Expression profiling of trehalose metabolism genes revealed significant downregulation of *SfTPS/TPP* (**Fig. 4L**) and *SfTreh1* (**Fig. 4M**). Furthermore, a significant decrease in trehalase activity was observed in *SflncRNA1*_OE larvae (**Fig. 4N**). Metabolomics analysis indicated increased trehalose and reduced glucose accumulation in *SflncRNA1*_OE larvae (**Fig. 4O-R**). *lncRNA1* overexpression deregulates trehalose homeostasis by suppressing the expression and resultant enzymatic activity of key genes in *H. armigera* and *S. frugiperda*.

To further validate these findings in a cell-culture-based model, *S. frugiperda* ovarian cell line (Sf9) was transfected with a *HalncRNA1* construct (**Fig. 5A**). qRT-PCR confirmed the *HalncRNA1* overexpression and downregulation of its putative target genes, *SfTPS/TPP* and *SfTreh1* (**Fig. 5B-D**). *HalncRNA1_*OE cells showed decreased *Sf*TPP and *Sf*Treh1 enzymatic activities (**Fig. 5E, F**). Metabolomic estimation depicts reduced intracellular accumulation of glucose-6-phosphate, trehalose-6-phosphate, trehalose, and glucose (**Fig. 5G–J**). Together, these results demonstrate that *HalncRNA1* and *SflncRNA1* act as negative regulators of trehalose metabolism, underscoring a conserved lncRNA-mediated mechanism controlling energy homeostasis in Lepidopteran insects.

**Fig 5.**
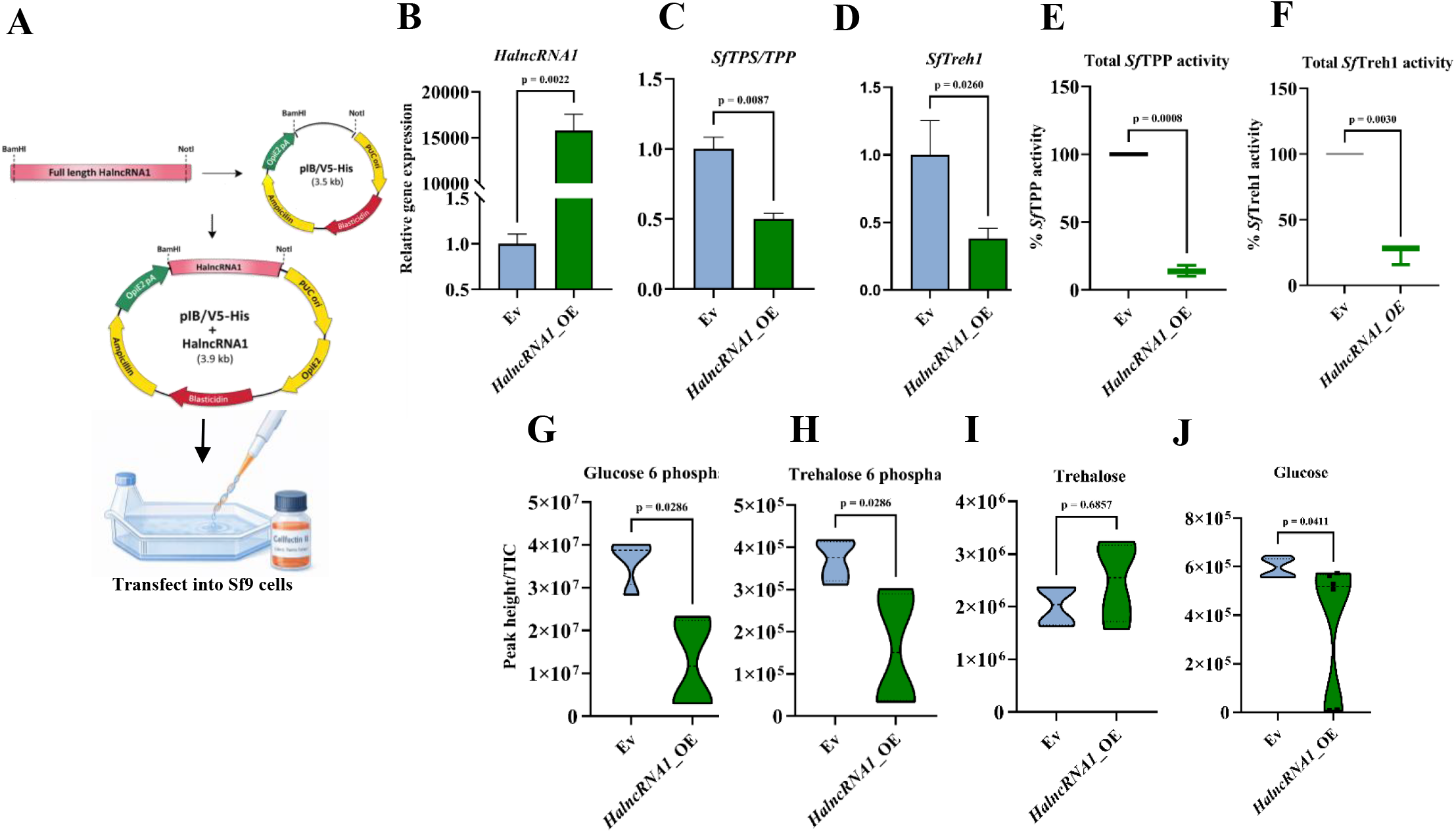
*lncRNA1* overexpression suppresses trehalose biosynthesis and degradation pathways in Sf9 cells. **(A)** Schematic representation of cloning of full-length *HalncRNA1* into the pIB/V5-His expression vector and transfection into Sf9 cells using Cellfectin II reagent. **(B)** qRT-PCR validation of *HalncRNA1* overexpression (OE) compared to empty vector (Ev) control. **(C, D)** Relative expression levels of *SfTPS/TPP* and *SfTreh1* following *HalncRNA1* overexpression in Sf9 cells. **(E, F)** Corresponding enzyme activity assays showing reduced *Ha*TPP and *Ha*Treh1 activity upon overexpression. **(G–J)** Metabolite profiling showing levels of glucose-6-phosphate, trehalose-6-phosphate, trehalose, and glucose in control and *HalncRNA1*-overexpressing cells. Differences between Ev and *HalncRNA1* OE treatments were assessed using an unpaired two-tailed Student’s *t*-test. Data are presented as mean ± SEM from three biological replicates. The exact *P*-value is indicated in the figure, with *P* < 0.05 considered statistically significant.

### *SflncRNA1* forms molecular associations with *TPS/TPP* and *Treh* mRNAs

To elucidate the molecular interactome of *SflncRNA1*, crosslinked RNA pull-down assays were performed using biotinylated antisense probes specific to *SflncRNA1* (**Fig. 6A**). Successful overexpression of *SflncRNA1* was first confirmed by qRT-PCR **(Fig. S4A)**. Semi-quantitative PCR analysis of the recovered RNA revealed enrichment of *SflncRNA1* (**Fig. 6B**) and its putative target *SfTPS/TPP* (**Fig. 6C**) and *SfTreh1* (**Fig. 6D**) transcripts in the *SflncRNA1* pull-down fraction. These transcripts were undetectable in the LacZ probe control, demonstrating the high specificity of the interaction. The recovery of *TPS/TPP* and *Treh1* mRNAs indicates that *SflncRNA1* engages in potential molecular interactions in a conserved manner with key components of the trehalose metabolic genes.

**Fig 6.**
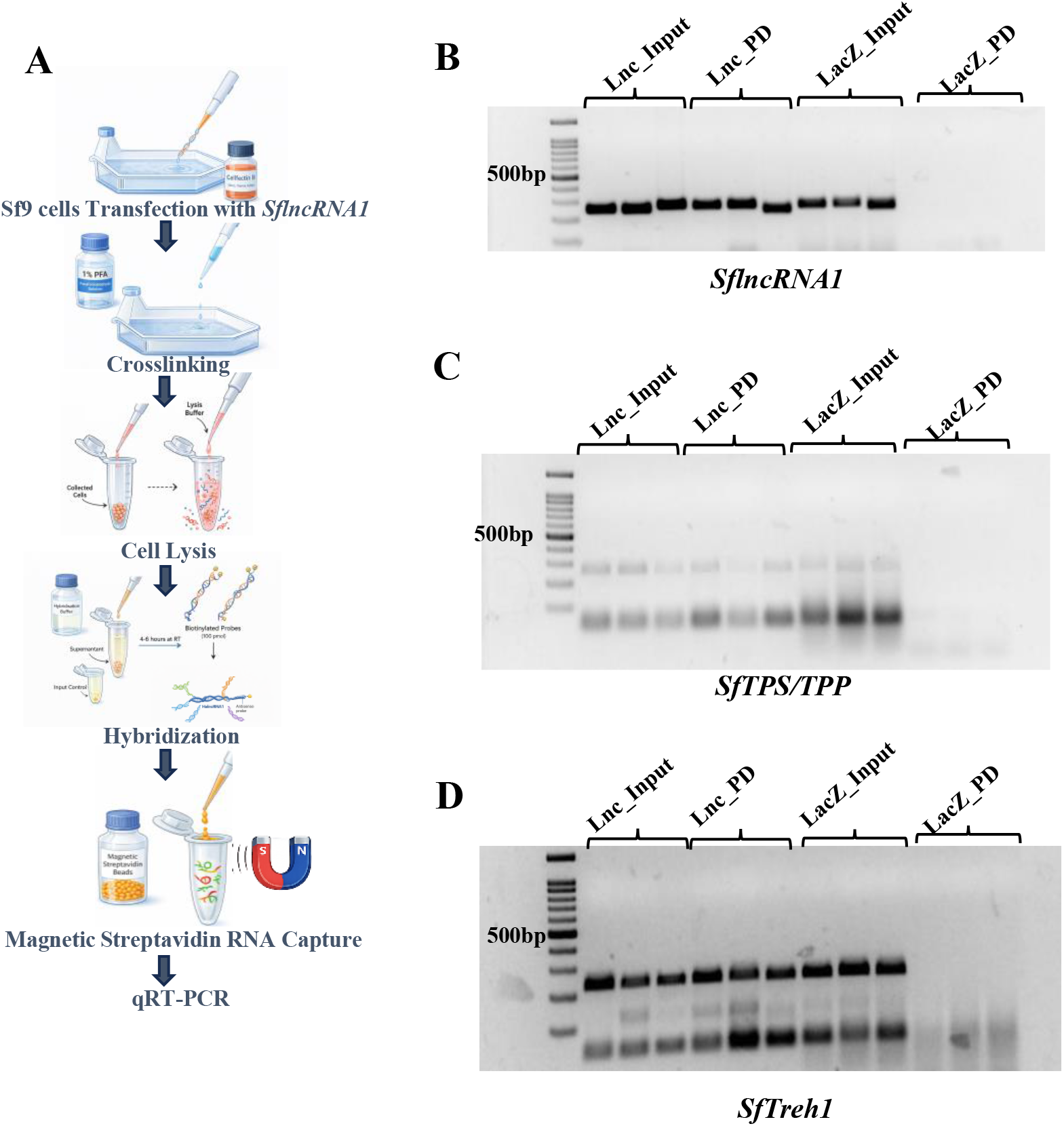
*SflncRNA1* forms molecular associations with *TPS/TPP* and *Treh* mRNAs. **(A)** Schematic workflow of the RNA pull-down assay in Sf9 cells. Cells transfected with *SflncRNA1* were crosslinked, lysed, and incubated with biotinylated antisense probes targeting *lncRNA1*, followed by capture using magnetic streptavidin beads. The enriched RNA complexes were subsequently analyzed by qRT-PCR. **(B–D)** Agarose gel electrophoresis showing enrichment of *SflncRNA1* **(B)**, *SfTPS/TPP* **(C)**, and *SfTreh1* **(D)** transcripts in pull-down (PD) samples compared to input controls, with LacZ probe serving as a negative control.

### Evolutionary relationship between *lncRNA1* and trehalose metabolism genes across lepidopteran insects showed high conservation

Phylogenetic reconstruction of *lncRNA1* (**Fig. 7A**), *TPS/TPP* (**Fig. 7B**), and *Treh* (**Fig. 7C**), sequences revealed pronounced order-level conservation across Insecta. The *lncRNA1* phylogeny resolved into well-supported, lineage-specific clades, including a strongly monophyletic assemblage of noctuid Lepidoptera (moths and butterflies) and hymenopteran (bees, wasps and ants)-specific clusters. Similarly, *TPS/TPP* and *Treh* phylogenies exhibited robust monophyly within Lepidoptera, with moths forming highly supported clades (bootstrap ≥99) and butterflies emerging as sister groups. Collectively, these results demonstrate that *lncRNA1*, *TPS/TPP* and *Treh* transcripts are evolutionarily conserved within their respective insect orders, highlighting the presence of order-specific regulatory and metabolic architectures preserved through lineage diversification. BLAST-based retrieval of homologous ncRNA sequences **(Supplementary Data 1F)** followed by Clustal Omega alignment revealed multiple highly conserved nucleotide blocks within *lncRNA1* across Lepidopteran species (**Fig. 7D**). RNA-RNA interaction models showed stable duplexes between lncRNAs and target mRNAs in insects, with minimum free energy (ΔG) values indicating strong binding. *SflncRNA1*–*SfTPS/TPP*, displayed an extended helical interaction with ΔG = −19.34 kcal/mol, featuring continuous base-pairing in the red-highlighted *SflncRNA1* region (**Fig. 7E**). *SflncRNA1-SfTreh* RNA interaction exhibited a similar elongated stem-loop structure with ΔG = −11.45 kcal/mol, suggesting high intramolecular or pairing stability (**Fig. 7F**). *HalncRNA1–HaTPS/TPP*, reveals a highly stable, multi-helical duplex with ΔG = −12.97 kcal/mol, the strongest among the pairs (**Fig. 7G**). *HalncRNA1–HaTreh*, showed a less extensive helical interaction with ΔG = −12.88 kcal/mol, shows favorable but weaker than the *HaTPS/TPP* targets (**Fig. 7H**). These predictions support potential lncRNA-mediated regulation of trehalose metabolism genes and merit experimental validation.

**Fig 7.**
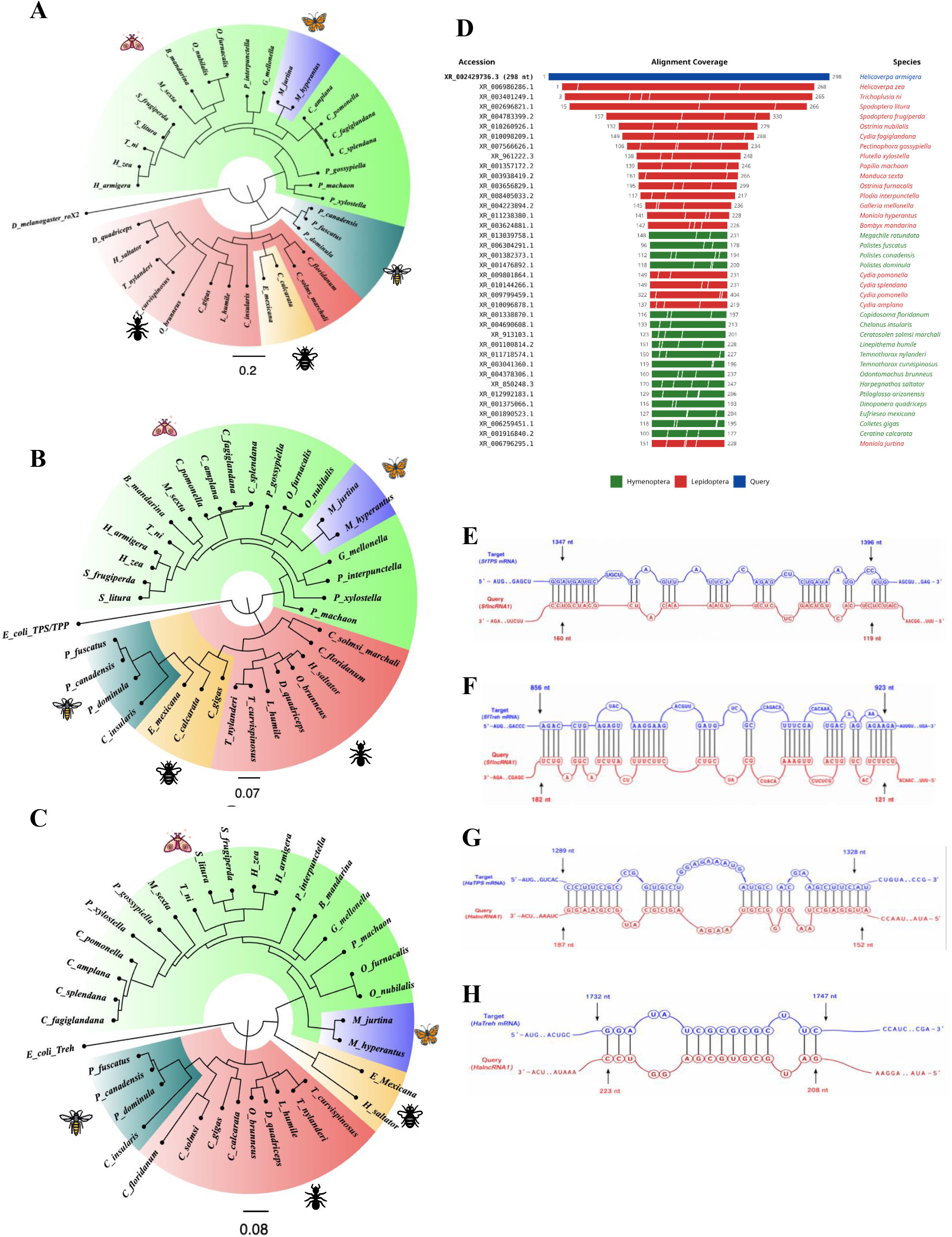
Evolutionary relationship between *lncRNA1* and trehalose metabolism genes across lepidopteran insects showed high conservation. **(A)** Phylogenetic tree illustrating the evolutionary relationships of *lncRNA1* across representative insect species, with conservation observed within major insect orders; *D. melanogaster roX2* lncRNA was used as the outgroup. **(B)** Phylogenetic reconstruction of trehalose-6-phosphate synthase/phosphatase (*TPS/TPP*) mRNA sequences across insect taxa, rooted with *Escherichia coli OtsA* (designated as *TPS/TPP* in the phylogeny). **(C)** Phylogenetic reconstruction of *Treh* mRNA sequences across insect species, *E. coli treA* (designated as *Treh* in the phylogeny). **(D)** Alignment coverage profile of *lncRNA1* homologues across multiple insect species, indicating conserved and variable regions together with sequence identity distribution across taxa. **(E)** Predicted interaction between *SflncRNA1* and *SfTPS/TPP* mRNA. **(F)** Predicted interaction between *SflncRNA1* and *SfTreh* mRNA. **(G)** Predicted interaction between *HalncRNA1* and *HaTPS/TPP* mRNA. **(H)** Predicted interaction between *HalncRNA1* and *HaTreh* mRNA. RNA–RNA interaction structures were predicted using computational interaction modeling. Nucleotides highlighted in yellow represent lncRNA1 regions involved in base-pairing interactions, whereas the corresponding target mRNA nucleotides are shown in black. The extensive complementary pairing observed across all four interactions supports a conserved lncRNA1-mediated post-transcriptional regulatory mechanism targeting key trehalose metabolism genes in both *H. armigera* and *S. frugiperda*.

Conserved sequence blocks identified within *lncRNA1* were further assessed for RNA-RNA interaction potential using the IntaRNA server. Comparative analysis revealed that *TPS/TPP* and *Treh* mRNAs from multiple Lepidopteran species exhibited strong predicted binding to their respective endogenous *lncRNA1* sequences **(Supplementary Data 1G)**. Notably, interaction sites consistently mapped to the conserved region spanning positions around 120 nt and extending to nearly 230 nt in *lncRNA1* **(Supplementary Data 1H)**. These conservation patterns indicate that evolutionarily conserved motif’s likely function RNA interaction domains.

### Deletion of conserved RNA interaction motifs disrupts *lncRNA1*-mediated regulation of trehalose metabolism

To confirm involvement of conserved motif in RNA interaction and potential regulation of trehalose metabolism, we created deletion mutants of *HalncRNA1* and *SflncRNA1* (**Fig. 8A**). *SflncRNA1Δ*^118–160^ (*SfTPS/TPP* interacting region deletion mutant), *SflncRNA1*Δ^120–182^ (*SfTreh* interacting region deletion mutant), and *HalncRNA1Δ^207-224^ (HaTreh* interacting region deletion mutant) were generated using extension overlap PCR and confirmed by qRT-PCR and gel electrophoresis **(Fig. S3A-B)**. Overexpression of *SflncRNA1* wild type (WT) (**Fig. 8B**) resulted in a significant downregulation of *SfTPS/TPP* and *SfTreh* transcript levels (**Fig. 8C, D**). While, *SflncRNA1Δ^118-160^* mutant overexpression (**Fig. 8E**), did not show any significant changes in *SfTPS/TPP* or *SfTreh1* expression compared to control (**Fig. 8F, G**).

**Fig 8.**
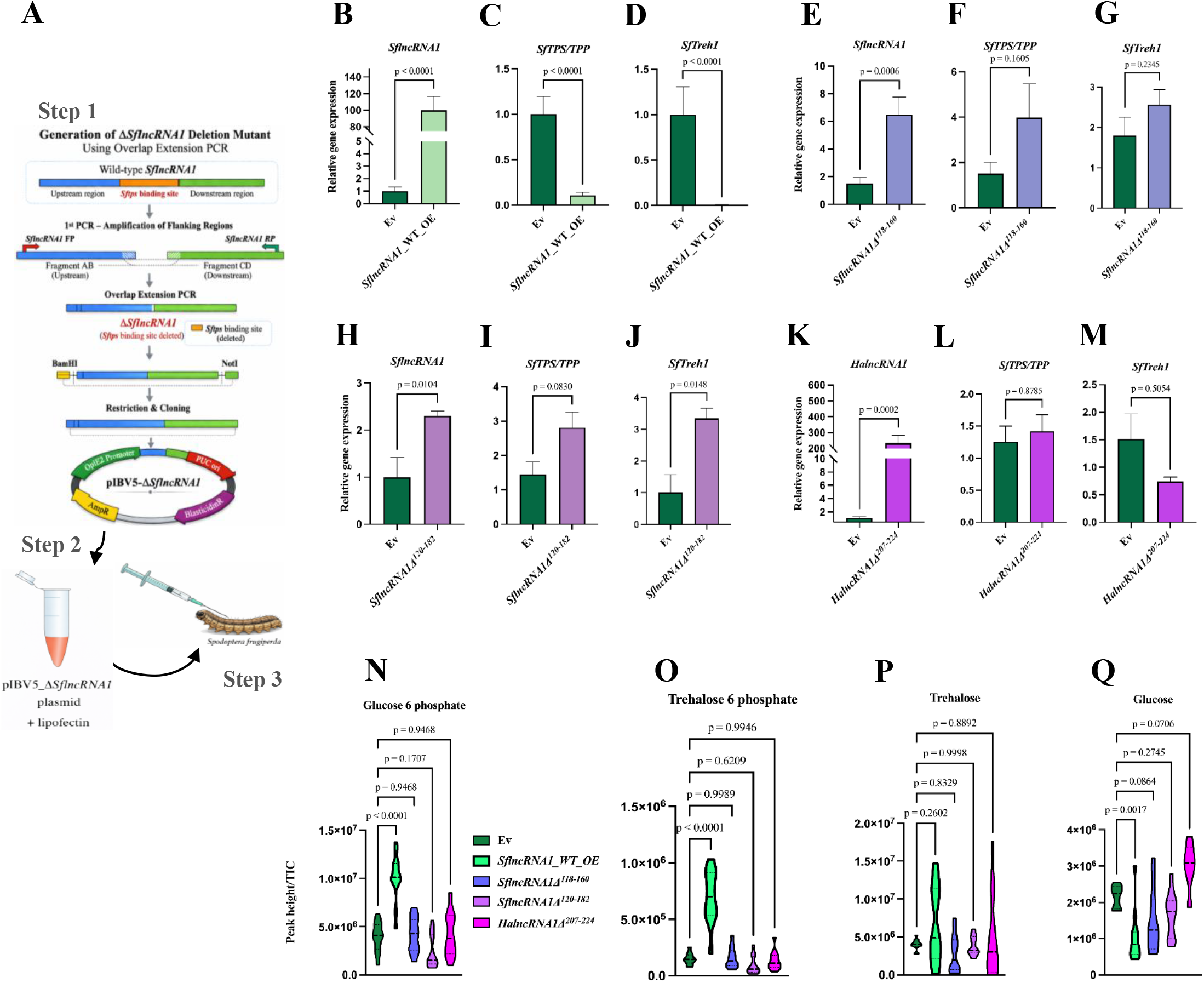
Deletion of conserved RNA interaction motifs disrupts *lncRNA1*-mediated regulation of trehalose metabolism. **(A)** Schematic representation of the generation of the Δ*SflncRNA1* deletion mutant lacking the predicted *SfTPS/TPP*-binding region using overlap extension PCR and cloning into the pIB/V5-His vector, followed by delivery into larvae. **(B–D)** Overexpression of wild-type *SflncRNA1* (WT_OE) results in significant downregulation of *SfTPS/TPP* and *SfTreh1* expression compared to empty vector (Ev) control. **(E–G)** Overexpression of *SflncRNA1Δ^118-160^* (*SfTPS/TPP* interacting region deletion mutant) shows increased *SflncRNA1Δ^118-160^* expression but fails to significantly alter *SfTPS/TPP* and *SfTreh1* transcript levels significantly. **(H–J)** Overexpression of the *SflncRNA1Δ^120-182^* (*SfTreh* interacting region deletion mutant) similarly shows elevated *SflncRNA1Δ^120-182^* expression without significant changes in *SfTPS/TPP* or *SfTreh1* expression. **(K–M)** Overexpression of the corresponding *HalncRNA1Δ^207-224^ (HaTreh* interacting region deletion mutant) shows elevated *HalncRNA1Δ^207-224^* expression with no significant changes in target gene expression. Differences between Ev and *SflncRNA1* (WT_OE), *SflncRNA1Δ^118-160^, SflncRNA1Δ^120-182^* and *HalncRNA1Δ^207-224^* mutants were assessed using an unpaired two-tailed Student’s *t*-test. Data are presented as mean ± SEM from three biological replicates. The exact *P*-value is indicated in the figure, with *P* < 0.05 considered statistically significant. Quantification of glucose-6-phosphate **(N)**, trehalose-6-phosphate **(O)**, trehalose **(P)**, and glucose **(Q)** levels in insects expressing wild-type *SflncRNA1* (WT)_OE with deletion mutants of *SflncRNA1Δ^118-160^* (*SfTPS/TPP* interacting region deletion mutant), *SflncRNA1Δ^120-182^* (*SfTreh* interacting region deletion mutant) and *HalncRNA1Δ^207-224^ (HaTreh* interacting region deletion mutant) compared to empty vector (Ev) control. Differences between Ev and *SflncRNA1* (WT_OE), *SflncRNA1Δ^118-160^, SflncRNA1Δ^120-182^* and *HalncRNA1Δ^207-224^* mutants metabolites were assessed using an unpaired two-tailed Student’s *t*-test. Data are presented as mean ± SEM from three biological replicates. The exact *P*-value is indicated in the figure, with *P* < 0.05 considered statistically significant.

Similarly, *SflncRNA1Δ^120-182^* mutant overexpression (**Fig. 8H**), did not alter the expression of target genes (**Fig. 8I, J**). Furthermore, overexpression of the *HalncRNA1Δ^207-224^* mutant (**Fig. 8K**) also did not alter the expression pattern of target genes (**Fig. 8L, M**). In addition, metabolite profiling revealed that overexpression of SflncRNA1_WT significantly increased the levels of glucose-6-phosphate, trehalose-6-phosphate, and trehalose, while reducing glucose levels. In contrast, overexpression of the *SflncRNA1* and *HalncRNA1* deletion mutant did not result in any detectable changes in these metabolites (**Fig. 8N–Q**). All these observations support that conserved RNA-RNA interaction motifs within *lncRNA1* are essential for regulating trehalose metabolism and maintaining downstream metabolite homeostasis.

### *SflncRNA1* modulates the post-transcriptional dynamics of trehalose metabolism-related transcripts in Sf9 cells

To investigate the potential post-transcriptional regulatory role of *SflncRNA1* in trehalose metabolism, an Actinomycin D-mediated transcript decay assay was performed in EGFP-overexpressing (EGFP-OE) and *SflncRNA1*-overexpressing (*SflncRNA1*-OE) Sf9 cells. Successful overexpression of *SflncRNA1* was confirmed by qRT-PCR (**Fig. 9A**). Following transcriptional inhibition with Actinomycin D, the transcript abundance of *SfTPS/TPP*, *SfTreh*, and *SflncRNA1* was quantified at 0, 2, 4, and 8 hour to assess their temporal expression dynamics (**Fig. 9B**). No visible phenotypic alterations were observed in Sf9 cells during the 8-hour Actinomycin D treatment period **(Fig. S6A)**. Temporal expression analysis of *SfTPS/TPP* revealed significant effects of time (p = 0.0004), treatment (p = 0.0016), and the time × treatment interaction (p = 0.0001), indicating that *SflncRNA1* overexpression significantly influenced the temporal expression dynamics of *SfTPS/TPP* under transcriptionally arrested conditions (**Fig. 9C and Fig. S6B)**. In EGFP-OE cells, *SfTPS/TPP* transcript abundance progressively increased following Actinomycin D treatment, whereas transcript levels remained consistently lower in *SflncRNA1*-OE cells throughout the time course. These findings suggest that *SflncRNA1* modulates the post-transcriptional regulation and turnover dynamics of *SfTPS/TPP* transcripts.

**Fig 9.**
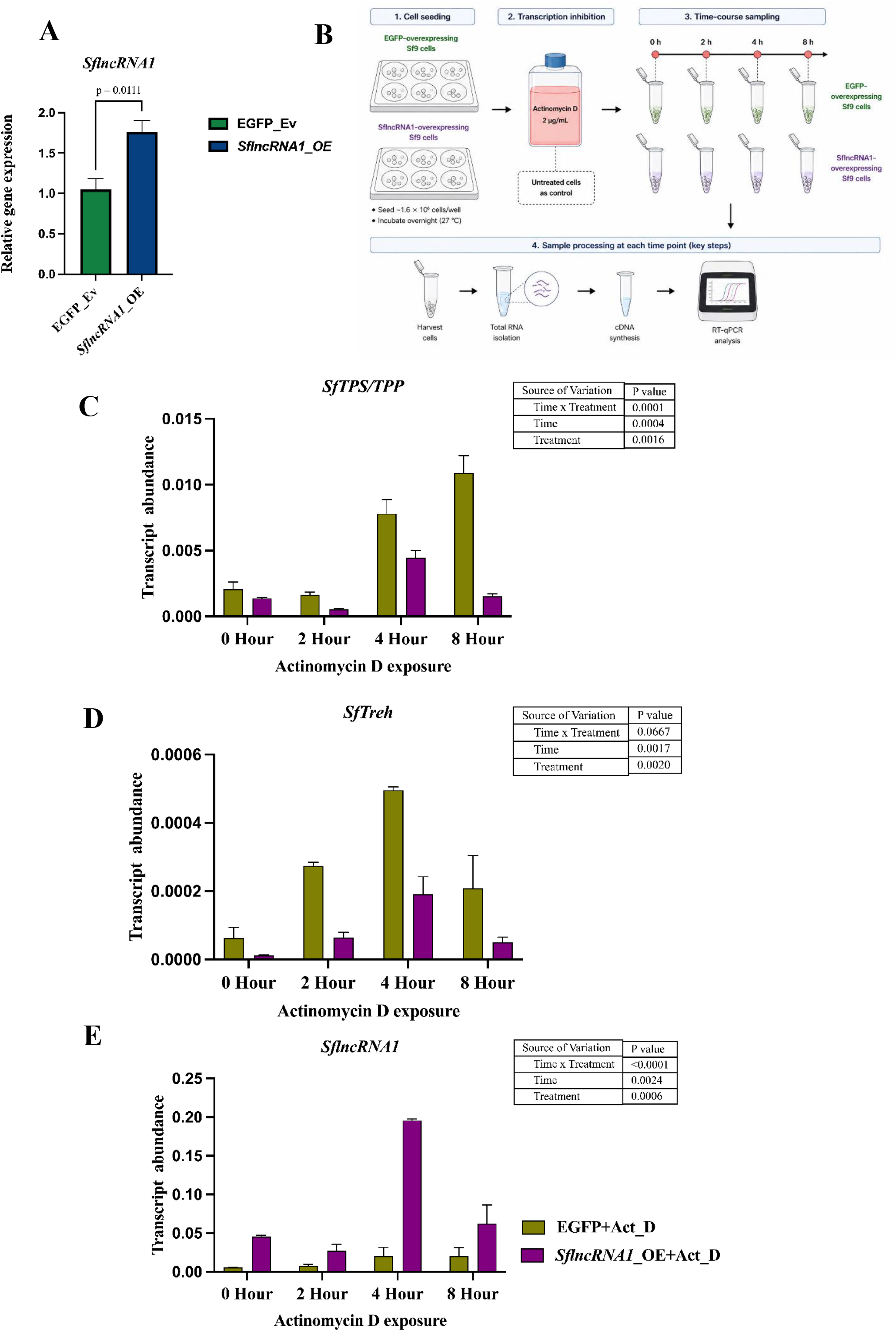
*SflncRNA1* modulates the post-transcriptional dynamics of trehalose metabolism-related transcripts in Sf9 cells. **(A)** Validation of *SflncRNA1* overexpression in Sf9 cells by qRT-PCR. **(B)** Schematic representation of the Actinomycin D-based transcript decay assay. EGFP-overexpressing and *SflncRNA1*-overexpressing Sf9 cells were treated with Actinomycin D (2 μg mL⁻¹), and samples were collected at 0, 2, 4, and 8 hour for RNA isolation and RT-qPCR analysis. (C–E) Temporal changes in transcript abundance of *SfTPS/TPP* **(C)**, *SfTreh* **(D)**, and *SflncRNA1* **(E)** following transcriptional inhibition. Transcript levels were quantified by qRT-PCR at the indicated time points after Actinomycin D treatment. Statistical significance of the effects of time, treatment, and their interaction was determined by two-way ANOVA, and corresponding P values are shown in the inset tables. Data represent mean ± SE of three biological replicates.

Similarly, temporal expression analysis of *SfTreh* revealed significant effects of time (p = 0.0017) and treatment (p = 0.0020), whereas the time × treatment interaction exhibited a marginally non-significant trend (p = 0.0667), indicating that both Actinomycin D exposure and *SflncRNA1* overexpression influenced *SfTreh* transcript dynamics (**Fig. 9D and Fig. S6C)**. Following transcriptional inhibition, *SfTreh* transcript abundance increased transiently over time, with comparatively lower transcript levels observed in *SflncRNA1*-OE cells than in EGFP-OE controls, suggesting a potential modulatory effect of *SflncRNA1* on *SfTreh* transcript turnover. In contrast, *SflncRNA1* transcript abundance showed highly significant effects of time (p = 0.0024), treatment (p = 0.0006), and the time × treatment interaction (p < 0.0001), confirming successful overexpression and dynamic transcript behavior during transcriptional inhibition (**Fig. 9E and Fig. S6D)**. Notably, *SflncRNA1*-OE cells consistently exhibited substantially higher transcript abundance than EGFP-OE controls across all time points. Collectively, these findings demonstrate that *SflncRNA1* significantly modulates the temporal transcript dynamics of key trehalose metabolism-associated genes under Actinomycin D-mediated transcriptional arrest, supporting its potential role as a post-transcriptional regulator of trehalose metabolic pathways.

## Discussion

Spatio-temporal expression dynamics of lncRNAs plays key role in insect development, metabolism and stress response ^16,31^. This pattern is consistent with the enrichment of *HalncRNA1* in late larval (L5) and pupal stages, as well as in metabolically active tissues such as gut, fat body, and head, suggesting its regulatory role. In this study, we found that *HalncRNA1* showed largely reciprocal expression to *TPS/TPP* and *Treh1* across developmental stages and tissues. Along with epigenetic and transcriptional level regulation, lncRNAs also act post-transcriptionally by binding mRNAs and RNA-binding proteins to control mRNA stability, processing, localization, and translation ^32–34^. Numerous metabolic lncRNAs in mammals function as trans regulators of enzymes and transporters in glycolysis, lipid metabolism, and glucose homeostasis ^33–35^. In parallel, large-scale insect lncRNA atlases predict extensive lncRNA connectivity to glycolytic and TCA cycle enzymes, supporting a conserved role for lncRNAs in metabolic network control ^16,31^. Within this framework, *HalncRNA1* emerges as a nuclear, structurally stable long intergenic noncoding RNA (lincRNA) that potentially fine-tunes trehalose-glucose homeostasis in lepidopteran larvae.

The pronounced structural stability and predicted pseudoknots of *HalncRNA1* are consistent with well-characterized lncRNAs such as *MALAT1* and H19 having defined secondary and tertiary structures to scaffold regulatory complexes and modulate transcriptional programs or mRNA processing in metabolic tissues ^33–36^. *HalncRNA1* likely uses discrete structural elements to both bind trehalose-pathway mRNAs directly and interact with nuclear regulatory factors.

*lncRNA1* knockdown increases *TPS/TPP* and *Treh* transcript levels and enzymatic activities, reduces trehalose, and elevates glucose, consistent with enhanced trehalose catabolism. In contrast, transient overexpression in larvae and Sf9 cells suppresses TPS/TPP and Treh expression and activity and reduces trehalose and glucose, as well as their phosphorylated intermediates. This functional data supports a model in which *lncRNA1* acts as a conserved, dose-dependent repressor of both trehalose synthesis and degradation. Potentially *lncRNA1*-mediated regulation of trehalose metabolism may shape growth, chitin biosynthesis, and energy balance in Lepidoptera and other model insects ^9,37^.

Importantly, the effects of *HalncRNA1* extended beyond the core trehalose pathway and were accompanied by extensive transcriptomic and metabolomic reprogramming. Silencing of *HalncRNA1* induced genes involved in glycogen mobilization, glycolysis, mitochondrial metabolism, and energy turnover, including glycogen phosphorylase, pyruvate kinase, phosphoglyceromutase, ATP synthase subunit f, adenylate kinase 5, and D-erythronate dehydrogenase. These transcriptional changes were paralleled by elevated levels of glucose, glucose-6-phosphate, pyruvate, lactate, malate, fumarate, and α-ketoglutarate, indicating enhanced carbon flux through glycolytic and TCA-cycle pathways ^1,38^. Together, these findings suggest that lncRNA1-mediated regulation of trehalose metabolism influences broader carbohydrate utilization networks and serves as a critical determinant of energy allocation in developing larvae.

The metabolic reprogramming observed in HalncRNA1-silenced insects was further accompanied by activation of nutrient-sensing and anabolic pathways. Increased expression of AKT, Rheb, IGF2, eIF4E, FASN1, FASN3, FABP, and LpR1, together with accumulation of fatty acids and storage lipids, indicates a shift toward lipid biosynthesis, nutrient storage, and biomass production. Notably, although ATP levels were reduced, multiple genes associated with mitochondrial energy metabolism and adenine nucleotide turnover were induced, suggesting elevated energy demand and rapid ATP utilization rather than impaired energy generation. Similar molecular signatures have been linked to activation of insulin/IGF and TOR signaling pathways, which coordinate nutrient sensing, lipid metabolism, protein synthesis, and growth in insects^39–41^. This coordinated metabolic state provides a plausible mechanistic explanation for the increased body size and body weight observed following HalncRNA1 silencing.

Pull-down experiments and interaction modeling converge on a direct duplex-based mechanism. Crosslinked pulldowns using antisense probes against *SflncRNA1* enrich *TPS/TPP* and *Treh* mRNAs, while in silico predictions revealed long, energetically favorable helices-particularly between *SflncRNA1* and *TPS/TPP* transcripts, indicative of stable RNA-RNA pairing. Conserved sequence blocks within lncRNA1, retained across Lepidopteran species, precisely overlap the predicted interaction sites. Targeted deletion of these motifs in *SflncRNA1* abolishes their ability to repress *TPS/TPP* or *Treh* expression and to modulate trehalose and glucose levels. This modular, motif-dependent repression parallels other lncRNA-mRNA systems in which short structural domains encode target specificity and post-transcriptional silencing capacity ^34^.

Additional evidence for post-transcriptional regulation was obtained from Actinomycin D-mediated transcriptional arrest assays. Following inhibition of de novo transcription, overexpression of *SflncRNA1* significantly altered the temporal dynamics of *TPS/TPP* and *Treh* transcripts, resulting in persistently lower transcript abundance relative to control cells. These observations indicate that *SflncRNA1* influences the behavior of target transcripts after transcriptional arrest and provide independent functional support for a post-transcriptional regulatory mechanism. When considered together with the RNA pull-down data, these findings strongly support a model in which *lncRNA1* regulates trehalose metabolism through direct RNA–RNA interactions that modulate transcript dynamics.

Phylogenetic analyses further support a coevolutionary relationship between *lncRNA1* and trehalose-pathway genes. *lncRNA1*, *TPS/TPP* and *Treh* each form well-supported, order-specific clades within Insecta, with strong monophyly of Lepidopteran lineages and conserved sequence blocks within *lncRNA1* across moth species. These patterns are consistent with previous findings that, although most lncRNAs evolve rapidly, some retain short conserved motifs and similar expression patterns across species, often associated with essential physiological functions ^17,37,42^. In Lepidoptera and other insects, trehalose metabolism is indispensable for larval fitness, molting, cuticle formation, and stress tolerance ^9,37^. The observed co-conservation suggests that a Lepidoptera-specific *lncRNA1*–TPS/Treh regulatory axis has become integrated into the trehalose metabolic network, complementing the diversification of trehalases and trehalose transporters that regulate tissue-and stage-specific carbohydrate metabolism ^43^.

Despite these insights, several limitations should be considered. First, RNAi-based approaches cannot distinguish whether the observed effects arise from the RNA molecule itself or from transcriptional activity at the locus. Second, off-target and systemic effects of dsRNA may contribute to some of the broader transcriptomic changes. Third, although RNA-RNA interactions are supported by pull-down and modeling, direct mapping of binding sites and interacting proteins is lacking. Fourth, transcriptomic analyses were performed on whole insects, potentially obscuring tissue-specific regulatory roles. Finally, evolutionary conservation was examined in a limited number of species, and broader validation is needed to establish generality across insect lineages. In addition to these, several questions remain open in this study. The precise molecular consequence of lncRNA-mRNA interaction, whether it affects stability, translation, or localization, has not yet been defined. The quantitative relationship between *HalncRNA1* levels and metabolic flux is also unclear, particularly under physiological conditions. In addition, the downstream physiological effects of altered trehalose metabolism, including impacts on molting, chitin synthesis, and survival, remain to be systematically explored.

Future studies may therefore focus on resolving the post-transcriptional mechanism should be resolved by measuring mRNA stability and translation efficiency of *TPS/TPP* and *Treh* under gain-and loss-of-function conditions, coupled with the identification of associated RNA-binding proteins. Furthermore, tissue-and stage-specific roles need to be examined using spatial transcriptomics or targeted perturbation approaches to capture cell-type-specific regulation. Also, cross-species functional assays, including motif swapping and rescue experiments, will help determine the evolutionary constraints and modularity of *lncRNA1*-mediated regulation.

In conclusion, this study uncovers a conserved lncRNA-based regulatory layer controlling trehalose metabolism in insects. By integrating RNA structure, molecular interactions, transcriptomic and metabolomic reprogramming, and physiological outcomes, we demonstrate that *lncRNA1* functions as a key regulator of energy homeostasis, coordinating trehalose metabolism with broader processes including carbohydrate utilization, nutrient sensing, lipid biosynthesis, and growth. These findings provide a mechanistic framework for understanding how non-coding RNAs fine-tune metabolic networks during insect development and highlight lncRNA-mediated regulatory pathways as potential targets for sustainable pest management strategies.

## Methods

### lncRNA Identification and secondary structure stability analysis

ncRNA sequences of *H. armigera* were retrieved from the NCBI genome assembly GCF_030705265.1. A total of 1,733 annotated non-coding RNA transcripts were initially extracted from the genome dataset. From these, transcripts specifically annotated as lncRNAs were selected according to annotation criteria, yielding 1,663 high-confidence lncRNA sequences that were used for subsequent structural analyses. To evaluate their intrinsic thermodynamic stability, we performed their secondary structure prediction locally using the ViennaRNA package (version 2.5.1) ^44^ with the RNAfold program. Partition function folding was enabled to compute ensemble thermodynamic properties using the command: RNAfold -p -d2 --noLP. The-p option calculated base-pairing probabilities and ensemble statistics, -d2 applied standard dangling-end energy parameters, and --noLP prevented isolated base pairs from forming unstable helices. For each lncRNA transcript, the following structural parameters were extracted: Minimum Free Energy (MFE; kcal/mol), Ensemble Free Energy (kcal/mol), Centroid Free Energy (kcal/mol), MFE frequency (%), and Ensemble Diversity. Because RNA thermodynamic values are inherently length-dependent, raw energy values were normalized by transcript length (in nucleotides) to eliminate size bias and enable cross-transcript comparisons.

The following normalized metrics were calculated: MFE_norm (MFE divided by sequence length; kcal/mol/nt), Ensemble_norm (ensemble free energy per nucleotide), Centroid_norm (centroid free energy per nucleotide), and Diversity_norm (ensemble diversity normalized per nucleotide). Additionally, Energy_gap was computed as the difference between Ensemble Free Energy and MFE, representing the thermodynamic separation between the optimal structure and the ensemble average.

To classify structurally stable lncRNAs, a two-dimensional stability framework was constructed by plotting MFE_norm against Diversity_norm. This integrative approach allows simultaneous evaluation of thermodynamic strength and ensemble compactness. Transcripts exhibiting highly negative MFE_norm values together with low Diversity_norm values clustered in the lower-left quadrant of the plot and were interpreted as structurally stable candidates. Cutoff thresholds of ≤ −0.2 kcal/mol/nt for MFE_norm and ≤ 0.2 per nucleotide for Diversity_norm were applied to define strong thermodynamic stability and low conformational variability, respectively. lncRNAs that satisfied both criteria were considered to possess stable and dominant secondary structures.

### Multimodal deep learning validation of non-coding potential

Following structural stability screening, coding potential assessment was performed to further validate the non-coding nature of the selected lncRNA candidates. For this purpose, the alignment-free deep learning framework lncRNA_Mdeep ^45^ was employed as a multimodal deep learning predictor for distinguishing lncRNAs from protein-coding transcripts. Structurally stable lncRNAs identified from thermodynamic stability analysis were evaluated for coding potential. Each transcript was provided as input to lncRNA_Mdeep, which generated a probability score ranging between 0 and 1. The default decision boundary of 0.5 was applied, with transcripts having a prediction probability ≥ 0.5 were classified as non-coding, and those with probability < 0.5 as protein-coding. This additional validation step ensured that the selected candidates possessed both thermodynamically stable secondary structures and strong computational support for non-coding potential.

### Pseudoknot prediction and structural refinement

Pseudoknot prediction was performed using IPknot++ (Version 2.3.1) ^46^, which predicts maximum expected accuracy (MEA) RNA secondary structures including pseudoknots based on base-pairing probabilities. To address length-dependent computational complexity, two models were applied: lncRNAs ≤500 nt were analyzed using the McCaskill partition function with ViennaRNA parameters, while lncRNAs >500 nt were processed using the LinearPartition-V (lpv) model with ViennaRNA parameters for efficient approximation of partition functions in long sequences.

In the IPknot++ output, pseudoknot interactions are denoted by square brackets (“[” and “]”). The total number of pseudoknot base pairs per transcript was determined by counting the number of opening brackets (“[”), each representing one pseudoknot interaction. To eliminate transcript length bias, pseudoknot density was calculated as the number of pseudoknot base pairs divided by transcript length (nt), providing a normalized measure of structural enrichment. To evaluate the differences in predicted structural features, we utilized the Wilcoxon Rank-Sum test (Mann–Whitney U test) to compare the distributions of pseudoknot counts and densities. Statistical significance was determined at p<0.05. Only structurally stable, coding-validated lncRNAs predicted to contain pseudoknot interactions were retained for downstream analyses. Subcellular localization of *HalncRNA1* was predicted using the lncLocator tool ^47^.

### Insect culture

*H. armigera* and *S. frugiperda* eggs were obtained from the ICAR–National Bureau of Agricultural Insect Resources (ICAR-NBAIR), Bengaluru, India. After hatching, larvae were reared on a chickpea-based artificial diet (AD) ^48^ at 28 °C, 80% relative humidity (RH), and a 16:8 h light:dark photoperiod. Second-generation insects were used for feeding and ‘-omics’ experiments, while third instar larvae were used for overexpression assays.

### Trehalose pathway inhibition and differential transcriptome analysis in *H. armigera*

*H. armigera* insects were fed with Validamycin A (VA), a known inhibitor of trehalase (*Ha*Treh-1) ^49^, and N-(phenylthio) phthalimide (NPP), identified as an inhibitor of TPP enzyme activity ^29,50^. Following inhibition of trehalose metabolism, transcriptome analysis was performed to identify differentially expressed genes in VA-and NPP-treated insects.

### RNA extraction and quantitative reverse-transcriptase PCR (qRT-PCR) analysis

Total RNA was isolated from *H. armigera* and *S. frugiperda* developmental stages (1st, 3rd and 5th instar larvae, pupa, adult male and female), dissected tissues (hemolymph, gut, fat body, head and cuticle), and experimental samples (control, gene-silenced and overexpression groups) using RNAiso Plus reagent (TaKaRa, Shiga, Japan) according to the manufacturer’s instructions, followed by DNase treatment with RQ1 RNase-free DNase (Promega, Madison, USA) to remove genomic DNA contamination. First-strand cDNA was synthesized from 2 μg of total RNA using oligo(dT) primers and the High-Capacity cDNA Reverse Transcription Kit (Applied Biosystems, Foster City, USA). Quantitative real-time PCR (qRT-PCR) was performed using TB Green Premix Ex Taq II (TaKaRa, Shiga, Japan) on a 7500 Fast Real-Time PCR System (Applied Biosystems, Foster City, USA), with gene-specific primers designed using Primer-BLAST (NCBI) (**Supplementary Table 1C**). The thermal cycling conditions consisted of an initial denaturation at 95°C for 30 second, followed by 40 cycles of 95°C for 5 second and 55°C for 30 second, along with melt curve analysis to confirm specificity. Relative expression levels of *HalncRNA1*, *HaTPS/TPP* and *HaTreh1* across developmental stages and tissues were estimated as transcript abundance using the 2^(-ΔCt) method, whereas relative gene expression under experimental treatments were calculated using the 2^(-ΔΔCt) method. *H. armigera actin* (*HaAct3ab*) and *S. frugiperda elongation factor 1-alpha* (*SfEf1-α*) used as internal reference genes; all reactions were performed with three biological replicates.

### lncRNA1 silencing in H. armigera and S. frugiperda by RNAi

Partial fragments of *HalncRNA1* and *SflncRNA1* were cloned into the L4440 vector for RNAi using gene-specific primers, and the resulting constructs were confirmed by sequencing (**Supplementary Data 1A**). The resulting recombinant vectors (L4440-*HalncRNA1*; L4440-*SflncRNA1*), and the L4440 empty vector, were transformed into *E. coli* HT115 (DE3) cells. To prepare the bacteria expressing dsRNA, a single colony was inoculated into Luria-Bertani (LB) broth supplemented with 100 µg/mL ampicillin and 12.5 mg/mL tetracycline antibiotics and incubated for 14 hours at 37°C with agitation at 180 rpm. The culture was then diluted 100-fold in 200 mL yeast tryptone (2xYT) medium and grown at 37°C with agitation at 180 rpm, with appropriate antibiotics, until the optical density (OD) reached 0.4. The synthesis of dsRNAs was induced by adding 1 mM isopropyl β-D-1-thiogalactopyranoside (IPTG), and these cultures were incubated for 5 hours under similar conditions. Bacterial cells were then pelleted by centrifugation at 4000 rpm at 4°C. Confirmation of dsRNA expression was achieved by extracting total RNA from aliquots of induced and uninduced culture using TRIzol reagent (TaKaRa Bio, Shiga, Japan) according to the manufacturer’s instructions, and dsRNAs were visualized on a 2% agarose gel.

### Feeding bioassay of *HalncRNA1-*dsRNA and *SflncRNA1*-dsRNA expressing bacteria to *H. armigera* and *S. frugiperda*

Bacterial cultures of dsRNA targeting *HalncRNA1*, *SflncRNA1* and L4440 empty vector were used for feeding second instar *H. armigera* and *S. frugiperda* larvae respectively. IPTG-induced 200 mL bacterial cultures were centrifuged at 4000 rpm for 10 minutes at 4°C, then resuspended in 4 mL of DEPC water. AD was prepared according to the methodology described by Chikate et al. ^48^. Bacterial cultures were then added to the AD at a final optical density (OD) of 1. Feeding bioassays were conducted on second instar larvae of *H. armigera* and *S. frugiperda*, with 30 larvae per treatment. The larvae were fed either an artificial diet containing either an L4440 empty vector bacterial culture (Control) or bacterial cultures expressing L4440-*HalncRNA1*-dsRNA or L4440-*SflncRNA1*-dsRNA. Insect growth and survival on both the control and dsRNA-containing diets were monitored on alternate days until pupation. On the 6th day, 20 larvae from each group were harvested by freezing them in liquid nitrogen. These larvae were then pooled into groups of five and treated as a single replicate. Silencing efficiency of *HalncRNA1* and *SflncRNA1* was evaluated by qRT-PCR. For transcriptomic analysis, 100 mg of *H. armigera* tissue exhibiting *HalncRNA1* silencing was collected per sample and submitted for RNA sequencing, with three biological replicates. In parallel, insects treated with the L4440 empty vector (EV) were used as controls, also in biological triplicates.

### RNA-seq data processing, alignment, and differential expression analysis

The raw fastq reads were pre-processed using Fastp v.0.23.4 (parameters: --trim_poly_g –cut front_--cut_tail --average_qual 30) ^51^. The rRNA reads were filtered from the trimmed fastq files using RiboDetector v0.3.1 ^52^ (parameters: -e rrna). The rRNA filtered reads were aligned to STAR-indexed *H. armigera* (NCBI, GCF_030705265.1) using the STAR aligner v 2.7.9a ^53^. The rRNA and tRNA features were removed from the GTF file of the *H. armigera*. The alignment file (sorted BAM) from individual samples were quantified using featureCounts v. 2.0.1 ^54^ based on the filtered GTF file to obtain gene counts. Differential expression estimation was done based on the gene counts using DESeq2 ^55^.

To identify genes differentially expressed following *HalncRNA1* silencing, normalized RNA-seq read counts from ds*HalncRNA1* and control (L4440) samples were subjected to differential expression analysis. For each gene, the log₂ fold change (Log₂FC) and corresponding P-value were calculated. A volcano plot was generated by plotting Log₂FC on the x-axis against the −log₁₀(P-value) on the y-axis, allowing simultaneous visualization of expression magnitude and statistical significance. Genes passing the predefined fold-change and significance thresholds were classified as significantly upregulated or downregulated and were further examined to identify biological pathways associated with the *HalncRNA1*-silencing phenotype. Significantly differentially expressed genes were subjected to Gene Ontology (GO) and KEGG enrichment analysis using ShinyGO ^56^, with *H. armigera* genomes used as reference model, a significance threshold of 0.05, and False Discovery Rate (FDR) correction.

To gain mechanistic insights into the altered glucose and trehalose levels and increased body weight observed following *HalncRNA1* silencing, the expression patterns of key genes involved in glucose and glycogen metabolism were examined using RNA-seq data. Normalized expression values of selected genes were extracted, Z-score transformed, and visualized as a heatmap using the pheatmap package in R. Hierarchical clustering of genes and samples was performed using Euclidean distance and complete linkage. Red and blue colors indicate relatively high and low transcript abundance, respectively, enabling visualization of metabolic transcriptional reprogramming induced by *HalncRNA1* silencing. To investigate the molecular mechanisms underlying the increased body weight observed following *HalncRNA1* silencing, genes associated with nutrient sensing, growth regulation, lipid biosynthesis, and hormonal signaling pathways were examined using RNA-seq data. Normalized expression values of selected genes were extracted and grouped into three functional categories: nutrient sensing and growth signaling, lipid biosynthesis and anabolic metabolism, and hormonal/developmental regulation. Mean normalized expression values were visualized as bar plots to compare transcript abundance between control and ds*HalncRNA1*-treated insects. This targeted analysis enabled the identification of key regulatory pathways associated with enhanced growth, nutrient utilization, and lipid accumulation induced by *HalncRNA1* silencing. To investigate the metabolic basis of the increased body weight phenotype associated with *HalncRNA1* silencing, metabolite profiles related to carbohydrate metabolism, glycolysis, the TCA cycle, lipogenesis, amino acid metabolism, energy metabolism, and hormonal regulation were analyzed. Mean metabolite abundances from control and ds*HalncRNA1*-treated insects were normalized using Z-score transformation and visualized as a bubble plot, where bubble color represents relative metabolite abundance and bubble size corresponds to metabolite concentration. This targeted metabolomic analysis enabled the identification of metabolic pathways altered by *HalncRNA1* silencing and provided insights into the redistribution of carbon and energy resources associated with enhanced growth and body weight gain.

### Overexpression of HalncRNA1 and SflncRNA1 in H. armigera and S. frugiperda

To establish transient overexpression, full-length *HalncRNA1* and *SflncRNA1* were amplified using gene-specific primers (**Supplementary Data 1B**) and subsequently cloned into the pIB/V5-His-TOPO vector via TOPO cloning, followed by transformation into *E. coli* TOP10 cells.The pIBV5-*HalncRNA1* and pIBV5-*SflncRNA1* plasmids were then combined with lipofectamine (1:1 v/v) (Thermo Fisher Scientific, Waltham, USA) and incubated at room temperature for 30 minutes. Following this, third instar larvae of *H. armigera* and *S. frugiperda* were anesthetized on ice, and each larva was injected with 25 ng/mg body weight of the plasmid–lipofectin mixture into the abdomen, with 30 larvae per treatment group. Control larvae were injected with 25 ng/mg of pIBV5-EGFP plasmid. After injection, larvae were maintained on an artificial diet for 48 hour, after which they were collected and immediately frozen in liquid nitrogen. The samples were subsequently used for gene expression analysis by qRT-PCR.

### *HalncRNA1* overexpression in Sf9 cells by transfection

*HalncRNA1* was cloned into the pIBV5 vector for overexpression studies in Sf9 cells. Initially, 8 x 10^5 Sf9 cells were seeded in a T25 cell culture flask with 5 mL of SF900 II serum-free media (Thermo Fisher Scientific, Waltham, USA) and incubated for 12 hours at 27°C. Cellfectin II (Thermo Fisher Scientific, Waltham, USA) was then diluted with 100 µL of SF900 II media and incubated for 30 minutes at room temperature. Similarly, 3 µg of the pIBV5-*HalncRNA1* plasmid and 3 µg of the pIBV5-EGFP plasmid (control) were separately diluted in 100 µL of SF900 II media and incubated for 30 minutes at room temperature. After incubation, the Cellfectin and plasmids were mixed and incubated for an additional 30 minutes at room temperature to form a DNA lipid complex. The used Sf900 II medium was removed from Sf9 cells, and the DNA–lipid complex was added dropwise, followed by the addition of 3 mL of antibiotic-free Sf900 II medium. Cells were incubated at 27 °C for 12 hours. After incubation, the medium was replaced with fresh Sf900 II medium, and cells were further incubated for 24 hours. Subsequently, the medium was replaced with Sf900 II medium supplemented with 50 µg/mL blasticidin (HiMedia, Mumbai, India) for antibiotic selection, and cells were incubated for 72 hours at 27 °C. Following this period, cells were passaged in Sf900 II medium containing 25 µg/mL blasticidin and maintained at 27 °C for an additional 72 hours. Finally, cells were harvested for RNA isolation to validate *HalncRNA1* overexpression in Sf9 cells.

### *SflncRNA1* overexpression, crosslinking, and RNA pull-down in *S. frugiperda* (Sf9) cells

Sf9 cells were cultured in a T25 flask at a density of 4×10^4 cells in Sf900 II SFM (Thermo Fisher Scientific, Waltham, USA) cell culture medium and incubated at 27°C for 24 hours. Cells were transfected with a *SflncRNA1* expression construct using Cellfectin II reagent (Thermo Fisher Scientific, Waltham, USA), according to the manufacturer’s instructions. Overexpression of *SflncRNA1* was confirmed by qRT-PCR. RNA pull-down assays were performed following the protocol described by Torres et al. ^57^ with minor modifications. For crosslinking, the culture medium was aspirated, and cells were rinsed with 10 mL of phosphate-buffered saline (PBS, pH 7.4), then incubated with 10 mL of 1% freshly prepared paraformaldehyde (PFA) in PBS for 10 minutes at room temperature with gentle agitation. Crosslinking was quenched by adding 1 mL of 1.25 M glycine, and the cells were agitated for an additional 5 minutes. After aspirating the solution, cells were washed twice with 10 mL of PBS, scraped into 1 mL PBS, transferred to 1.5 ml microcentrifuge tubes, and centrifuged at 510 × *g* for 5 minutes at 4°C. The supernatant was removed, and the pellet was stored at –80°C. For RNA pull-down, thawed cell pellets were lysed by adding 1 mL of ice-cold lysis buffer per 100 mg of pellet, vortexed for 5 minutes, and centrifuged at 1,200 × *g* for 5 minutes at 4°C. Supernatants were collected, mixed with two volumes of hybridization buffer, and vortexed. From this, a 20 µL aliquot was saved as an input control. Biotinylated antisense oligonucleotide probes (100 pmol) **(Supplementary Data 1D)** targeting *SflncRNA1* were added, and samples were incubated for 4–6 hours at room temperature under moderate agitation. Then, 50 µL of pre-washed magnetic streptavidin beads supplemented with 200 U/mL RNase inhibitor and 5 µL/mL protease inhibitor cocktail were added, and the mixture was incubated overnight at room temperature on a tube rotator. The following day, samples were placed on a magnetic stand, and the supernatant was discarded. The beads were washed three times with 900 µL of wash buffer by gentle rotation for 5 minutes at room temperature, followed by magnetic separation. RNA was then extracted directly from the beads using the TRIzol reagent method, quantified using a NanoDrop spectrophotometer, and reverse-transcribed into cDNA using a reverse transcription kit according to the manufacturer’s instructions. The enrichment of pull-down targets was subsequently assessed by qRT-PCR.

### Metabolite extraction and LC–MS analysis

Each biological replicate consisted of five pooled insects (n = 5) from control and treated larvae for metabolite extraction. Three biological replicates were analysed. For metabolite extraction, 100 mg of crushed tissue was homogenized in 80% methanol (v/v) at a ratio of 1:5 (w/v). The suspension was vortexed for 30 seconds, bath-sonicated for 30 minutes at room temperature, and then centrifuged at 14,000g for 20 minutes. The supernatant was stored at −80°C overnight to allow protein precipitation. Subsequently, the supernatant was filtered through a 0.22 μm syringe filter, and the filtrate was transferred to sample vials. The samples were analyzed using an LC-HRMS (Q Exactive Orbitrap, Thermo Fisher Scientific, Waltham, USA) mass spectrometer with an electrospray ionization (±ESI) source for ionization. For LC-based metabolite separation, a Hypersil GOLD C18-25002-152130 column (2.1 × 150 mm, 1.9 μm particle size, Thermo Fisher Scientific, Waltham, USA) was employed, with the column temperature maintained at 35 °C and a flow rate of 0.3 ml/minute. Metabolites were separated using a 20-minute gradient with 100% MS grade water (Solvent A) and 100% MS grade acetonitrile (Solvent B), both containing 0.1% formic acid. The LC method commenced with 2% B for the initial 0.3 minutes and increased to 30% over the next 2 minutes. The B% was increased from 30% to 45% for 7 minutes, then raised to 98% for 12 minutes, after which it was held for the subsequent 3 minutes. The column was then equilibrated to the initial solvent ratio (98% A: 2% B) for the final 5 minutes. For peak extraction, baseline filtering, calibration, peak alignment, deconvolution, peak identification, and integration, MS-DIAL software version 4.9.221218 (http://prime.psc.riken.jp/Metabolomics_Software/MS-DIAL/index.html) was utilized, with an LC–MS/MS spectral database sourced from MassBank of North America. Quantification of metabolites involved in glucose and energy metabolism was carried out. In this study, LC-HRMS peak intensities were normalized using a total peak height (or area) / total ion current (TIC) normalization approach. Briefly, normalization was performed at the sample level by dividing the intensity of each detected feature by the sum of all peak intensities within the same sample. This procedure scales each feature relative to the total signal contribution of that sample, correcting for variation in sample loading, ionization efficiency, and instrument response across injections ^58^. The following formula was used for normalization:

Normalized peak intensity = Peak height (or area) / Total ion current (TIC)

### Trehalose 6-phosphate phosphatase activity

The enzymatic activity of *Ha*TPP was measured by monitoring the release of Pi from trehalose 6 phosphate (T6P) using a malachite green reagent (Sigma–Aldrich, St.Louis, USA) ^59,60^. In a final volume of 100 μL, the assay contained 1 mM T6P (Sigma–Aldrich, St.Louis, USA), 2 mM MgCl2 (Hi-Media, Thane, India), 50 mM phosphate buffer (pH 7), and the crude enzyme (50 μg). The reaction mixture was incubated for 30 minutes at 37°C in a water bath. Subsequently, the reaction was stopped by adding two volumes of a filtered solution containing 0.15% malachite green, 1% ammonium molybdate (Hi-media, Thane, India), and 12.5% (v/v) concentrated HCl (Thomas Baker, Mumbai, India). The mixture was left at room temperature for 5–7 minutes for colour development. The absorbance of the reaction mixture was measured at 630 nm.

### α, α-trehalase activity

The activity of total trehalase was determined by monitoring the release of glucose, a reducing sugar, from α, α-trehalose using the DNSA (Dinitrosalicylic acid) reagent (Sigma–Aldrich, St.Louis, USA) ^61^. In this assay, 50 µg of proteins were incubated with 150 μL of trehalose (0.25%) at 37°C for 15 minutes. Following the incubation period, 500 μL of DNSA reagent was added to stop the reaction. The reaction tubes were then placed in a boiling water bath for 5 minutes, and the absorbance was measured at 540 nm. One unit of trehalase activity was defined as the amount of enzyme required to release 1 μM of glucose per minute at 37°C under the given assay conditions.

### Evolutionary analyses of *lncRNA1*, *TPS/TPP*, and *Treh* across Insecta

To investigate the evolutionary relationships of *lncRNA1* across insect species, the *HalncRNA1* nucleotide sequence was queried against the NCBI core nucleotide (Core_nt) database using BLAST. All annotated ncRNA homologs were retrieved and used for phylogenetic tree construction. Similarly, for the same set of species, homologous *TPS/TPP* and *Treh* mRNA sequences were collected for comparative analysis. For phylogenetic analysis of *lncRNA1*, *D. melanogaster roX2* lncRNA was used as an outgroup. For phylogenetic analysis, *Escherichia coli OtsA* (designated as *TPS/TPP* in the phylogeny) was used as the outgroup for *TPS/TPP*, whereas *E. coli treA* (designated as *Treh* in the phylogeny) served as the outgroup for *Treh*. Multiple sequence alignments were generated using Muscle with default nucleotide substitution models appropriate for each dataset using MEGA 12.1 ^62^. Phylogenetic trees were constructed using the Maximum likelihood method implemented in W-IQ_TREE ^63^, and branch support was evaluated using 1,000 bootstrap replicates. Final tree topologies were visualized and annotated to highlight lineage-specific clustering patterns and evolutionary divergence across insect taxa using *FigTree v1.4.4*.^64^.

### Sequence conservation and RNA–RNA interaction analysis

The *HalncRNA1* sequence was subjected to BLAST analysis against the NCBI RefSeq RNA database to identify homologous lncRNAs across insect species and to detect conserved regions among different taxonomic classes. Annotated ncRNA sequences from the identified species were retrieved **(Supplementary Data 1F)**, and conserved regions were visualized in R using the *ggplot2* and *dplyr* packages ^65,66^. These conserved regions were subsequently analyzed for potential RNA–RNA interactions with species-specific *TPS/TPP* and *Treh* mRNA sequences using the IntaRNA server ^67^ with default parameters.

### Cloning of *HalncRNA1* deletion mutants

To investigate the functional interaction between *HalncRNA1*, *SflncRNA1* and their target mRNAs (*TPS/TPP* and *Treh*), putative binding regions were identified using the IntaRNA tool ^67^, which predicted conserved lncRNA–mRNA base-pairing sites. To assess the functional importance of these regions, deletion mutants of *SflncRNA1*Δ^118–160^ (*SfTPS/TPP* interacting region deletion mutant), *SflncRNA1*Δ^120–182^ (*SfTreh* interacting region deletion mutant), and *HalncRNA1Δ^207-224^ (HaTreh* interacting region deletion mutant) were generated using overlap extension PCR. Two primer pairs **(Supplementary Data 1E)** were designed to amplify the upstream and downstream flanking regions of the predicted binding sites. The primers were modified to include overlapping sequences, enabling fusion of the two fragments in a secondary PCR reaction and generating an internal deletion of the target region. The resulting PCR products were cloned into the pIB/V5 overexpression vector using enzyme digestion and ligation. Clones were validated by Sanger sequencing. Third instar larvae of *H. armigera* and *S. frugiperda* were anesthetized on ice, and each larva was injected with 25 ng/mg body weight of the plasmid–lipofectin mixture into the abdomen, with 30 larvae per treatment group. Larvae were injected with 25 ng/mg of pIB/V5-EGFP (negative control) or *SflncRNA1*_WT plasmid (positive control). After injection, larvae were maintained on an artificial diet for 48 hours, after which they were collected and immediately frozen in liquid nitrogen. Successful overexpression of the construct was confirmed by qRT-PCR.

### *SflncRNA1* modulates the post-transcriptional dynamics of trehalose metabolism-related transcripts in Sf9 cells

Approximately 1.6 × 10^6 EGFP-overexpressing and *SflncRNA1*-overexpressing Sf9 cells were seeded separately into each well of a 6-well plate and incubated overnight at 27°C. The following day, Actinomycin D (Sigma–Aldrich, St.Louis, USA) was added to the culture medium at a final concentration of 2 µg/mL in Sf-900 II medium to inhibit de novo transcription. Untreated EGFP-overexpressing and *SflncRNA1*-overexpressing Sf9 cells were maintained as controls. All treatments were performed in three biological replicates. Cells were harvested at four different time points following Actinomycin D treatment: 0 h, 2 h, 4 h, and 8 h. At each time point, cells were gently scraped, collected into 1.5 mL microcentrifuge tubes, and centrifuged to pellet the cells. The supernatant was discarded, and the cell pellets were washed once with 1X PBS (pH 7.5) followed by centrifugation again. The final cell pellets were immediately processed for total RNA isolation, followed by cDNA synthesis for downstream qRT-PCR analysis.

### Statistical analyses

All experiments were performed in triplicate, and data are presented as mean ± SEM from three biological replicates. Statistical analyses were conducted using GraphPad Prism v8.0 (GraphPad Software, San Diego, CA, USA). Differences between control and treatment groups were evaluated using an unpaired two-tailed Student’s *t*-test. Exact *P*-values are indicated in the figures, with *P* < 0.05 considered statistically significant.

## Data availability

RNA-seq data have been deposited in the NCBI Sequence Read Archive (SRA) under BioProject accession number PRJNA1446711.

## Acknowledgements

Vikram Nichit and Deepti Wagh acknowledges the University Grants Commission (UGC), India, for the junior and senior research fellowships. Anand Kumar Shukla acknowledges AcSIR, India, for the Integrated Dual-Degree Program (IDDP) and the research fellowship for IDDP under the CSIR-GATE-JRF program (Award No.: 31/ GATE/11(55)/2025-EMR-I). The authors would like to thank Dr. Vasudevan Seshadri and Gaurav Agarwal (NCCS, Pune) for their guidance in RNA pull-down experiments. The authors thank Dr. Kundan Sengupta (IISER, Pune) for generously providing Actinomycin D for use in this study. The authors thank Dr. Meenakshi Tellis for their suggestions and technical help. The authors sincerely thank Dr. Suvasri Dutta and Sharada Mohite for their critical review of the manuscript.

## Author contributions

V.N., and R.J. conceptualization; V.N., D.W., A.K.S., and R.J. methodology; V.N., D.W., and R.J. formal analysis; V.N., D.W. investigation; V.N., D.W., and R.J. visualization; V.N., A.K.S. *In-silco* data analysis; V.N.,R.J. writing – original draft; D.W., A.K.S., N.K., and R.J. writing – review and editing; R.J. resources; R.J. supervision, project administration, and funding acquisition.

## Supplementary figures legends

Fig S1. Silencing of *HalncRNA1* modulates the expression of trehalose metabolism, sugar transporter, and developmental regulatory genes in *H. armigera*. (A) Validation of *HalncRNA1* dsRNA production in HT115 cells carrying the recombinant L4440-HalncRNA1 construct. **(B-O)** Relative mRNA expression levels of genes associated with trehalose metabolism and related regulatory pathways were quantified by qRT–PCR in larvae fed with control dsRNA (L4440_Ev) or ds*HalncRNA1*. Genes analysed include (B) *HaTreh2*, sugar transporter genes (C) *HaST9*, (D) *HaST29*, (E) *HaST46*, and (F) *HaST64*, transcriptional and signalling regulators (G) *HaFOXO*, (H) *HaIGF2*, and (I) *HaAL*, and developmental regulators (J) *HaE2F5*, (K) *HaE2F7*, (L) *HaETS*, (M) *HaDFD*, (N) *HaFTZ*, and **(O)** *HaE93*. Differences between L4440 Ev and *HalncRNA1* dsRNA treatments were assessed using an unpaired two-tailed Student’s *t*-test. Data are presented as mean ± SEM from three biological replicates. The exact *P*-value is indicated in the figure, with *P* < 0.05 considered statistically significant.

Fig S2. *HalncRNA1* silencing reprograms trehalose and glucose metabolism, driving a shift in carbohydrate utilization pathways **(A)** Hierarchical clustering heatmap of the top 50 differentially expressed genes (DEGs) identified following *HalncRNA1* silencing. Rows represent genes and columns represent biological replicates of control (L4440) and ds*HalncRNA1*-treated insects. Gene expression values are shown as Z-scores, with red and blue indicating relatively high and low expression, respectively. The heatmap reveals distinct clustering of control and silenced samples, highlighting extensive transcriptomic reprogramming following *HalncRNA1* silencing. **(B)** Hierarchically clustered bubble heatmap of trehalose metabolism-related genes and selected sugar transporter transcripts in control (L4440) and ds*HalncRNA1*-treated insects. Bubble color represents Z-score normalized expression levels, with red indicating higher expression and blue indicating lower expression. The plot highlights distinct expression patterns of trehalose metabolic and transporter-associated genes following *HalncRNA1* silencing.

**Fig S3. *HalncRNA1* overexpression leads to significant downregulation of *HaTreh2* in *H. armigera*. (A)** Relative expression of *HaTreh2* was determined by qRT–PCR in larvae injected with pIBV5_EGFP empty vector (Ev) or *HalncRNA1* overexpression construct (HalncRNA1_OE). Differences between Ev and *HalncRNA1* OE treatments were assessed using an unpaired two-tailed Student’s *t*-test. Data are presented as mean ± SEM from three biological replicates. The exact *P*-value is indicated in the figure, with *P* < 0.05 considered statistically significant.

**Fig S4. (A)** Validation of *SflncRNA1* overexpression in Sf9 cells by qRT-PCR prior to RNA pull-down analysis. Relative transcript abundance of *SflncRNA1* was significantly higher in *SflncRNA1*-overexpressing (*SflncRNA1*-OE) cells compared with the EGFP-overexpressing control (EGFP-Ev), confirming successful overexpression. Data are presented as mean ± SEM from three biological replicates. The exact *P*-value is indicated in the figure, with *P* < 0.05 considered statistically significant.

**Fig S5. Molecular validation of *lncRNA1* deletion constructs by PCR. (A)** Agarose gel electrophoresis of PCR-amplified fragments corresponding to *SflncRNA1Δ^118-160^* and *SflncRNA1Δ^120-182^* deletion constructs alongside empty vector (Ev) control. **(B)** PCR verification of wild-type *SflncRNA1* (WT) and *HalncRNA1Δ^207-224^* deletion mutant. DNA size marker (left) indicates fragment size in base pairs (bp). Images are representative of independent clone validations.

**Fig S6. *SflncRNA1* modulates the post-transcriptional dynamics of trehalose metabolism-related transcripts in Sf9 cells. (A)** Representative bright-field images of EGFP-overexpressing (EGFP-OE) and *SflncRNA1*-overexpressing (*SflncRNA1*-OE) Sf9 cells following exposure to Actinomycin D for 0, 2, 4, and 8 h. No visible morphological alterations were observed during the treatment period. **(B–D)** Temporal expression profiles of *SfTPS/TPP* **(B)**, *SfTreh* **(C)**, and *SflncRNA1* **(D)** in EGFP-OE and *SflncRNA1*-OE cells with or without Actinomycin D treatment. Transcript abundance was quantified by qRT-PCR at the indicated time points. Statistical significance of the effects of time, treatment, and their interaction was determined by two-way ANOVA, and corresponding P values are shown in the inset tables. Data represent mean ± SE of three biological replicates.

## Supplementary Data legends

Supplementary Data 1A. Primers used for dsRNA synthesis of *lncRNA1*. Primer sequences used for amplification of *lncRNA1* fragments for dsRNA synthesis in *H. armigera* and *S. frugiperda*. Forward and reverse primers contain 5′ T7 promoter sequences to enable in vitro transcription.

Supplementary Data 1B. Primers used for amplification of full-length *lncRNA1* constructs. Primer sequences used to amplify full-length *lncRNA1* transcripts from *H. armigera* and *S. frugiperda* for cloning and overexpression assays.

**Supplementary Data 1C. Primers used for qRT–PCR and construct validation.** Primer sets used for quantitative gene expression analysis of *lncRNA1*, trehalose metabolism genes (*TPS/TPP*, *Treh1*, *Treh2*), sugar transporter genes, and reference genes. Primers used for vector confirmation and construct validation are also included.

**Supplementary Data 1D. Biotin-labelled probes used for RNA pull-down assays.** Sequences of 5′ biotin-labelled antisense oligonucleotides targeting *SflncRNA1*, along with LacZ control probes, used for RNA pull-down assays to identify interacting mRNAs.

Supplementary Data 1E. Primers used for generation of *lncRNA1* deletion mutants. Primer sequences designed to generate targeted deletions within predicted *lncRNA1* interaction regions with *TPS/TPP* and *Treh* transcripts in *H. armigera* and *S. frugiperda*.

Supplementary Data 1F. *lncRNA1* sequences used for multiple sequence alignment across Insecta. List of *lncRNA1* sequences from representative insect species, including accession IDs, used for comparative sequence analysis across Lepidoptera and Hymenoptera.

Supplementary Data 1G. IntaRNA-predicted interactions between *lncRNA1* and trehalose metabolism genes. Predicted *lncRNA1* interactions with *TPS/TPP* and *Treh* mRNAs across insect species using IntaRNA. Minimum free energy (kcal/mol), interacting nucleotide positions, and corresponding transcript accession IDs are provided.

Supplementary Data 1H. Predicted *lncRNA1* binding sites in *TPS/TPP* and *Treh* transcripts across Insecta. Sequences of predicted base-pairing regions between *lncRNA1* and *TPS*/*TPP and Treh* mRNAs identified using IntaRNA, indicating conserved interaction motifs involved in post-transcriptional regulation.

Supplementary Data 2. Computational identification and structural characterization of candidate lncRNAs in *Helicoverpa armigera*.

**Supplementary Data 3. Normalized expression profiles of growth-regulatory genes and metabolites associated with HalncRNA1 silencing.**

